# Detection of infiltrating fibroblasts by single-cell transcriptomics in human kidney allografts

**DOI:** 10.1101/2020.09.03.281733

**Authors:** Hemant Suryawanshi, Hua Yang, Michelle Lubetzky, Pavel Morozov, Mila Lagman, Gaurav Thareja, Alicia Alonso, Carol Li, Catherine Snopkowski, Aziz Belkadi, Franco B. Mueller, John R. Lee, Darshana M. Dadhania, Steven P. Salvatore, Surya V. Seshan, Vijay K. Sharma, Karsten Suhre, Manikkam Suthanthiran, Thomas Tuschl, Thangamani Muthukumar

## Abstract

We tested the hypothesis that single-cell RNA-sequencing (scRNA-seq) analysis of human kidney allograft biopsies will reveal distinct cell types and states and yield insights to decipher the complex heterogeneity of alloimmune injury. We selected 3 biopsies of kidney cortex from 3 individuals for scRNA-seq and processed them fresh using an identical protocol on the 10x Chromium platform; (i) HK: native kidney biopsy from a living donor, (ii) AK1: allograft kidney with transplant glomerulopathy, tubulointerstitial fibrosis, and worsening graft function, and (iii) AK2: allograft kidney after successful treatment of active antibody-mediated rejection. We did not study T-cell-mediated rejections. We generated 7217 high-quality single cell transcriptomes. Taking advantage of the recipient-donor sex mismatches revealed by X and Y chromosome autosomal gene expression, we determined that in AK1 with fibrosis, 42 months after transplantation, more than half of the kidney allograft fibroblasts were recipient-derived and therefore likely migratory and graft infiltrative, whereas in AK2 without fibrosis, 84 months after transplantation, most fibroblasts were donor-organ-derived. Furthermore, AK1 was enriched for tubular progenitor cells overexpressing profibrotic extracellular matrix genes. AK2, eight months after successful treatment of rejection, contained endothelial cells that expressed T-cell chemoattractant cytokines. In addition to these key findings, our analysis revealed unique cell types and states in the kidney. Altogether, single-cell transcriptomics yielded novel mechanistic insights, which could pave the way for individualizing the care of transplant recipients.

## Introduction

Molecular approaches complement conventional histopathology and have propelled precision transplantation medicine to the bedside [1–3]. Single-cell RNA-sequencing (scRNA-seq) provides hitherto unavailable opportunities to study cell types and cell states at an unprecedented level of precision [4–6]. Our goal was to investigate the utility of scRNA-seq at an individual patient level to address important conundrums in clinical transplantation. Given the complex heterogeneity of alloimmune rejection, we tested the hypothesis that single-cell transcriptomics—by enabling molecular phenotyping of the host infiltrating cells and donor parenchymal cells—will yield novel mechanistic insights, especially in the context of antibody-mediated injury, for individualizing the care of transplant recipients.

Immune rejection of the allograft remains a significant challenge despite the use of potent immunosuppressive drugs [7–9]. Rejection episodes restrict the benefits of transplantation and negatively impact long-term kidney allograft survival [10]. Treatment of rejection is constrained by the limited therapeutic armamentarium focused predominantly on the adaptive arm of the immune system and despite improvement in clinical and laboratory parameters, seldom achieves histological remission [10, 11]. Also, despite anti-rejection therapy, it is possible that allograft injury persists at a molecular level and perpetuates allograft dysfunction. It is tempting to speculate that effective treatment of the lingering immune injury may improve the long-term outcome of kidney transplant recipients. This, however, requires better understanding of the complex immune interactions between the recipient genome and the genome of the organ donor.

We studied two clinico-pathological scenarios: (i) chronic persistent tissue injury and worsening allograft function and (ii) resolved acute tissue injury following successful treatment of an episode of active antibody-mediated rejection. These results were compared to the single-cell transcriptomes of cells isolated from a native kidney used for living-donor kidney transplantation. We did not study T-cell-mediated rejection. We resolved 12 clusters of major cell types at the first level of single-cell gene expression analysis, with a subset of cell clusters further resolved by subclustering analysis. We identified 4 distinct fibroblast subpopulations differentially present in the biopsies and made the surprising finding that one fibroblast subtype in the transplant biopsies was kidney-recipient rather donor-derived. We also identified tubular progenitor cells with profibrotic gene signature. Finally, the transcriptomes of endothelial cell subtypes provided additional insights into the anti-allograft response.

## Materials and methods

### Tissue collection, dissociation, and single-cell preparation

We followed a standard operating procedure for performing kidney allograft biopsies to obtain samples for scRNA-seq. Tissue samples were collected under local anesthesia by real-time ultrasound guidance using an 18g Bard Monopty automated spring-loaded biopsy gun, and a Civco Ultra-Pro II in-plane needle guide attached to the ultrasound probe to prevent any contamination by tissues other than kidney. The presence of kidney cortical parenchyma without the presence of kidney medulla, kidney capsule, or any extra-renal tissue was verified by examination of the biopsy tissue under the microscope. The biopsies were subsequently transported in phosphate-buffered saline to our Gene Expression Monitoring laboratory and immediately dissociated for single-cell capture. We developed and used an in-house protocol for single-cell suspension preparation. In brief, the sample was placed in 400 μl of freshly prepared tissue dissociation solution comprised of 100 μl Liberase TL solution (2 mg/ml, Sigma-Aldrich), 500 μl Tyrode’s solution-HEPES-based (Boston BioProducts), and 200 μl DNase I solution (1 mg/ml, Stemcell technologies) and incubated at 37°C water bath for 15 min. The cell suspension was passed through a 40 μm Falcon™ cell strainer (ThermoFisher Scientific) into a 50 ml centrifuge tube filled with 5 ml fetal bovine serum (ThermoFisher Scientific), washed through with Dulbecco’s phosphate-buffered saline (ThermoFisher Scientific), and centrifuged for 5 min at 300g. Cells were resuspended in 2% bovine serum albumin (New England BioLabs) and were transferred immediately in ice to the genomics core laboratory.

### Single-cell capture, library preparation, sequencing, data processing, and generation of gene expression matrix

Single-cell suspension on ice was immediately transferred to the Weill Cornell Medicine genomics core facility. scRNA-seq libraries were prepared using the Chromium Single Cell 3’ Reagent Kit V2 (10x Genomics) according to the manufacturer’s instructions. The library was sequenced on Illumina HiSeq 2500 platform as follows: 26 bp (Read1) and 98 bp (Read2). The sequencing was performed to obtain 150–200 million reads (each for Read1 and Read2). The 10x raw data were processed individually for 3 kidney samples using previously described Drop-seq pipeline (Drop-seq core computational protocol V1.2, http://mccarrolllab.com/dropseq/) with the following parameters. The Read1 bases 1–16 was tagged with cell barcode ‘XC’ and bases 17–26 was tagged with a unique molecular identifier (UMI) ‘XM’. Low-quality bases containing reads were removed and the 3’-end of Read2 was trimmed to remove poly(A) sequences of six or more bases and were aligned to human genome (hg38) reference using STAR aligner (STAR_2.5.1a), allowing no more than three mismatches. The gene expression matrix was then generated using the ‘MIN_BC_READ_THRESHOLD= 2’ option to retain UMIs with read evidence of two or more.

### scRNA-seq data analysis

Cell clustering analysis was performed collectively for all 3 samples using ‘Seurat V3.1’, an R package for exploration of single-cell data. Briefly, only those genes that were expressed in more than three cells and cells that expressed more than 200 but less than 5000 genes were retained. Ubiquitously expressed genes such as ribosomal protein-coding (RPS and RPL) and non-coding RNA (*MALAT1*) genes were removed. We also removed miRNA and snoRNA genes from clustering analysis. The clustering analysis was performed in an iterative manner where in round 1 the cells with >25% mitochondrial content was identified and removed from round 2 clustering analysis. In round 2 analysis, mitochondrially coded genes (MT-) were also dropped from clustering analysis. We first generated three separate objects for AK1, AK2, and HK samples using ‘CreateSeuratObject’ function. Next, we merged these objects into one object using ‘merge’ function. We separated endothelial, epithelial, immune and stromal cells into their individual Seurat objects and conducted at least two rounds of analysis on each to identify and remove doublet captures. Typically, doublet capture show expression of markers of two different lineages (e.g., T cells and epithelial cells). Normalization was performed applying ‘logNormalize’ function of Seurat. Using function ‘FindVariableFeatures’ ∼2000 genes were identified, followed by ‘ScaleData’. Next, ‘RunPCA’, ‘FindNeighbors’, ‘FindClusters’, and ‘RunUMAP’ was used with different dimensions 1:10 and resolution of 0.5 wherever these options were needed. Reduction method “umap” was used and clusters were plotted using ‘DimPlot’ option. To identify differentially expressed genes by each cluster, Wilcoxon rank sum test inbuilt in the Seurat package was used with parameters ‘min.pct = 0.25’ and ‘thresh.use = 0.25’.

Subclustering analyses of the cell groups described in this manuscript were performed using similar strategy as described above. The expression of established lineage marker genes was used to assign cell types. Once the cell types were identified, average expression was calculated for each cell type followed by normalization to 10,000 to create a TPM (transcript per million)-like value.

For donor/recipient origin of cell analysis, we quantified the expression of female-specific gene (*XIST*) and male-specific Y chromosome autosomal genes (*RPS4Y, EIF1AY, DDX3Y*) as previously described [12–16]. We created separate Seurat objects for HK, AK1, and AK2 cells maintaining the original cell type identity. Using ‘DotPlot’ function, we plotted the expression of female-specific gene (XIST) and male-specific Y chromosome autosomal genes (*RPS4Y, EIF1AY, DDX3Y*) [12, 13, 15–19]. We verified the results of the donor/recipient origin of cell analysis by also clustering cells based on genotype. Raw sequencing data from each sample was individually aligned using 10x Genomics Cell Ranger 6.1.1 to human genome reference (GRCh38) downloaded from 10x Genomics (https://cf.10xgenomics.com/supp/cell-exp/refdata-gex-GRCh38-2020-A.tar.gz) using default parameters [20]. The output BAM and barcode file were used as inputs for ‘Singularity’ [21] version of ‘Souporcell’ pipeline [22]. The samples were individually processed with default parameters of pipeline with k = 2. The cluster file was further filtered to include only singlet cell cluster assignment. The cluster data was merged into Seurat object [23] for clustered fibroblast cells in R version 4.1.0 [24].

For cells of the peritubular capillaries (PTC), in order to perform differential gene expression analysis by samples, we could not employ the conventional differential gene expression analysis methods such as DESeq2 or edgeR, due to the small number of samples. For this analysis, we only allowed genes that had >2 TPM expression in at least one of the samples (HK, AK1, or AK2). Next, for each gene, we calculated the largest (maximal) and second largest TPM expression values across all three samples. Finally, only the genes with at least two-fold difference between maximal and second largest TPM were reported.

Normalized expression profiles of PT, PG1 and PG2 were generated and used for matrisome analysis. We utilized a previously described list of matrisome genes and then subset the list for each category of matrisomes (collagen, proteoglycan, and secreted factor) from the TPM expression profiles of PT, PG1, and PG2 cell types. Log_2_(TPM+1) values for top expressed genes in each of the matrisome category were represented in the heatmap.

To authenticate our annotation of cells, we used ‘Azimuth’ to map our data to an annotated reference dataset [23]. We mapped our data of 7,217 high quality cells from the three samples to the reference consisting of 64,693 kidney cells generated in the Human Biomolecular Atlas Program (HuBMAP) and the Kidney Precision Medicine Project (KPMP) [25].

### Study approval

The patients that we describe herein underwent living donor kidney transplantation and were followed up at our center—New York Presbyterian Hospital/Weill Cornell Medicine. Donor nephrectomies and recipient transplant surgeries were both done at our center and no organs or tissues were procured from individuals who were incarcerated. Patients provided written informed consent to participate in the study and the informed consent was obtained prior to their inclusion in the study. The research protocol was approved by the Weill Cornell Medicine Institutional Review Board (protocol number: 1404015008). The clinical and research activities that we report here are consistent with the principles of the “Declaration of Istanbul on Organ Trafficking and Transplant Tourism”[26]. This study did not generate new unique reagents. We have deposited our scRNA-seq data files, compliant with the single cell version of MIAME/MINSEQE standards [27], at NCBI’s Gene Expression Omnibus under the accession number GSE151671.

## Results

### Clinical characteristics and kidney cortical biopsy specimens

A summary of the clinical and histopathological characteristics of the healthy kidney donor and the two kidney transplant recipients is provided in Table 1. The two allograft biopsies included transplant glomerulopathy with graft dysfunction and follow up after active antibody-mediated rejection with normal graft function. We did not include biopsies from kidney allografts with chronic active T-cell-mediated rejection, interstitial fibrosis, and tubular atrophy not otherwise specified, or stable function and no acute rejection several years after transplantation. Briefly, biopsy HK was obtained from a healthy 40-year-old living kidney donor. The recipient who received kidney from this healthy living donor had immediate graft function and normal serum creatinine at 12 months after ––transplantation with no major infections or acute rejection. Biopsy HK was done at the operating room during the back-table preparation of the native kidney prior to its implantation in the recipient.

**Table 1.**
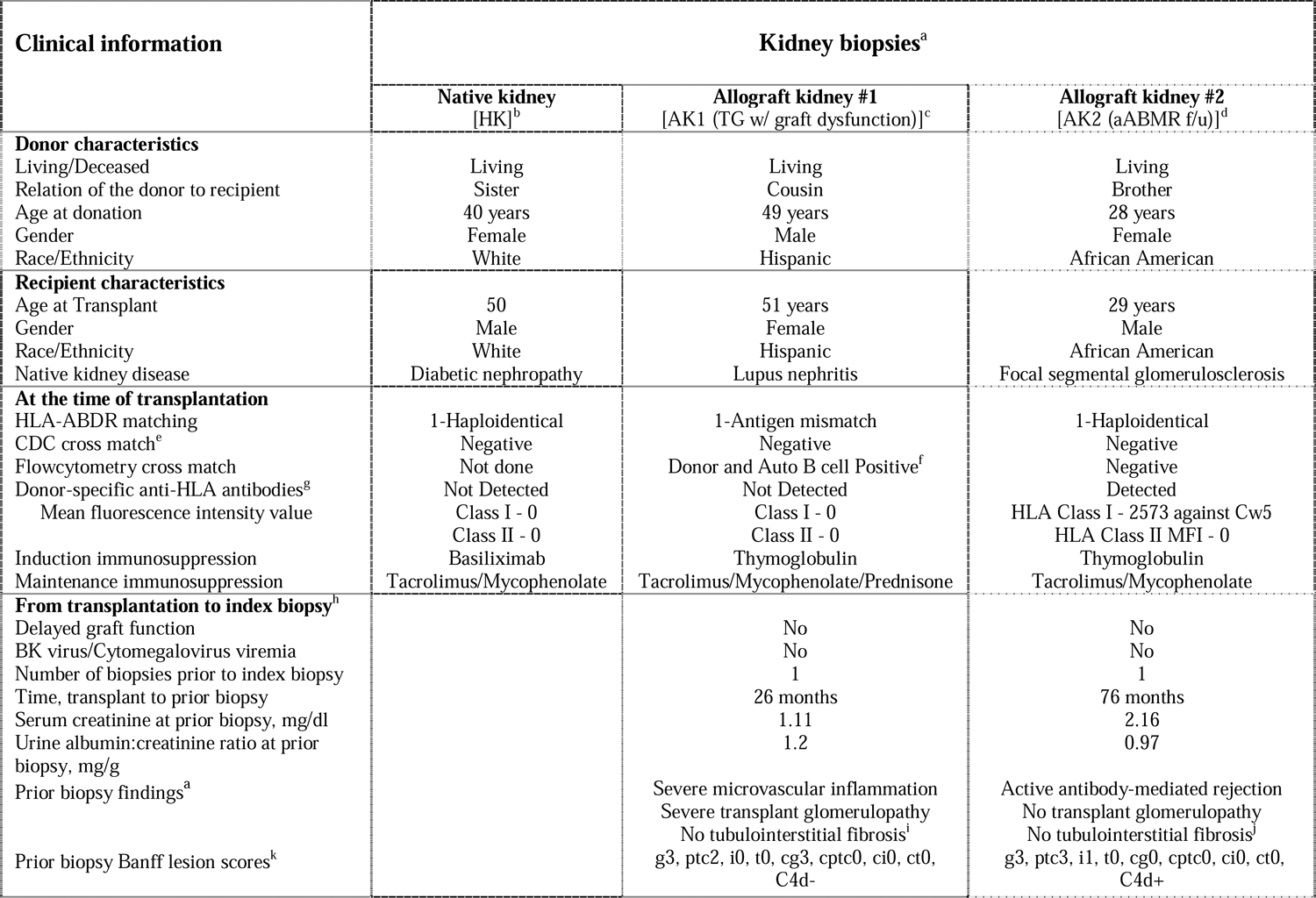
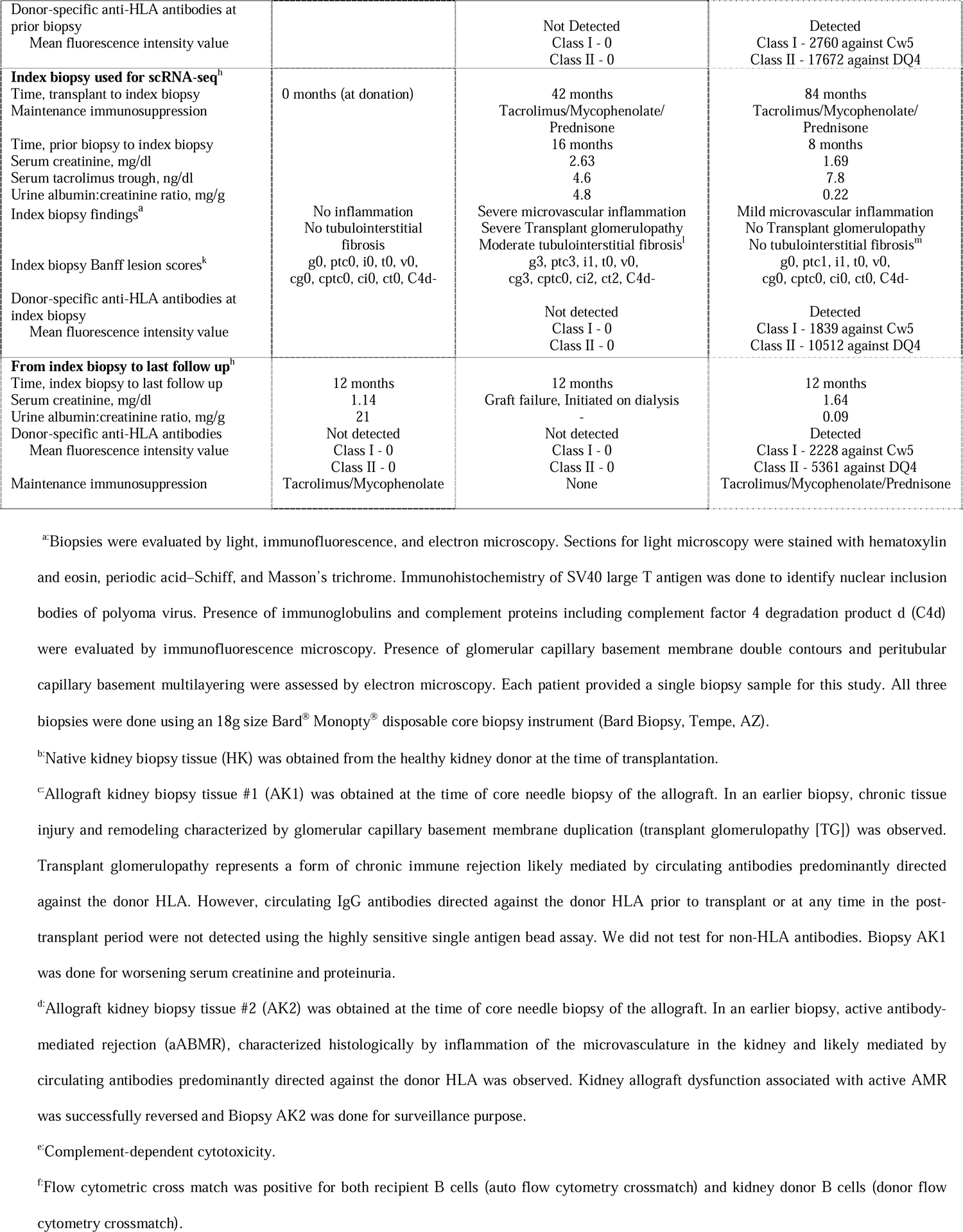
Clinical characteristics of the three subjects whose kidney biopsies were used for scRNA-seq

Biopsy AK1 was obtained by real-time ultrasound guidance from a 51-year-old woman. She developed end-stage kidney disease due to lupus nephritis and received a living-donor kidney transplant. She had worsening of proteinuria after the living-donor kidney transplantation and an allograft biopsy 26 months after transplantation revealed interstitial fibrosis and tubular atrophy, microvascular inflammation, and transplant glomerulopathy. The biopsy, however, did not fulfill the Banff criteria (an international standardized criteria for reporting allograft biopsies) for antibody-mediated rejection and there was no evidence for recurrence of lupus nephritis. Circulating immunoglobulin G antibodies directed against the donor HLA prior to transplant or at any time in the post-transplant period were not detected using the highly sensitive single-antigen bead assay. We did not test for non-HLA antibodies. Before transplant, she had a negative T and B cell complement-dependent cytotoxicity cross match and a negative T cell flowcytometry crossmatch. However, she had a positive B cell flowcytometry crossmatch to her kidney donor cells (donor flow cytometry B cell crossmatch) and to her own cells (auto flow cytometry B cell crossmatch). The index biopsy that was used for scRNA-seq was done for worsening proteinuria and kidney function 16 months after the initial biopsy (Fig 1 and S1 Fig).

**Fig 1.**
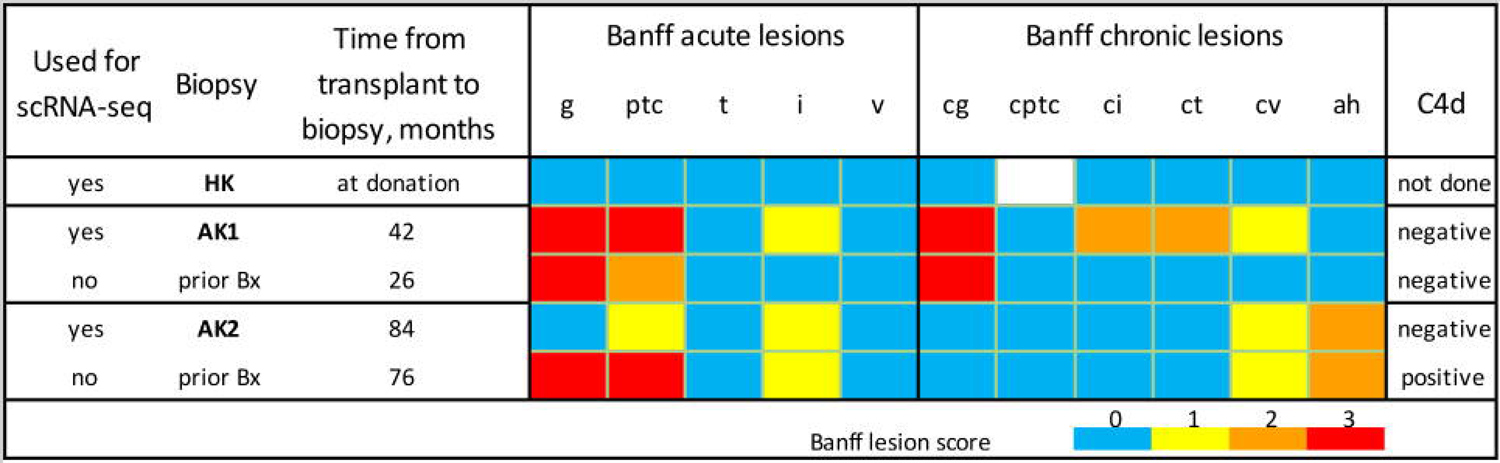
Histopathological characteristics of kidney biopsies Figure depicts the histopathological characteristics of the kidney biopsies used for scRNA-seq, HK, AK1 and AK2. Biopsy HK was obtained from a healthy 40-year-old living kidney donor and was done at the operating room during the back-table preparation of the native kidney prior to its implantation in the recipient. Kidney allograft biopsies AK1 was obtained by real-time ultrasound guidance from a 51-year-old woman. She had a prior biopsy that was done 16 months before the index biopsy used for scRNA-seq. Biopsy AK2 was obtained by real-time ultrasound guidance from a 29-year-old man. He had a prior biopsy that was done 8 months before the index biopsy used for scRNA-seq. Allograft biopsies, AK1 and AK2, were evaluated by light, immunofluorescence, and electron microscopy. Colors represent semi-quantitative Banff lesion scores from 0 to 3. Banff acute lesions include glomerular inflammation (g), peritubular capillary inflammation (ptc), tubular inflammation (t), interstitial inflammation (i), and vascular inflammation (v). Banff chronic lesions include chronic glomerulopathy (cg), peritubular capillary basement membrane multilayering (cptc), interstitial fibrosis (ci), tubular atrophy (ct), chronic vascular lesions (cv), and arteriolar hyaline thickening (ah). Peritubular capillary staining for complement split product 4d (C4d) by immunofluorescence was negative.

Subject AK2 had end stage kidney disease due to focal and segmental glomerulosclerosis. He developed acute elevation of serum creatinine, 76 months after a living-donor kidney transplantation and allograft biopsy revealed active antibody-mediated rejection (Banff category 2) with no chronic glomerular or tubulointerstitial changes. He was treated with our transplant center protocol comprised of methylprednisolone, plasmapheresis, intravenous immunoglobulin, and bortezomib and had resolution of graft dysfunction. The index biopsy that was used for scRNA-seq was done by the treating physician for surveillance purpose 8 months after the acute rejection episode (Fig 1 and S2 Fig). He had normal graft function (serum creatinine <2 mg/dl and albuminuria <500 mg/day) at the time of the index biopsy but had persistent circulating IgG antibodies directed against the donor HLA.

### Identification of distinct cell types in healthy and allograft kidneys

We conducted iterative cell clustering analysis using Seurat V3.1, an R package for exploration of single-cell data [28]. We obtained 9762 cells; 2545 cells with >25% mitochondrial content was removed from subsequent analysis. The final single-cell gene expression matrices for the three kidney biopsies were comprised of 7217 high-quality cells and separated into 12 cell clusters by gene expression (Fig 2A, left) with no contributions from batch processing (Fig 2A, right). Based on differential gene expression and previously established markers of cell types or states we designated these clusters as proximal tubular cells (PT1 and PT2), tubular progenitor cells (PG), loop of Henle cells, collecting duct cells, and intercalated cells (LH.CD.IC), fibroblasts (FB), endothelial cells (EC), pericytes and vascular smooth muscle cells (PC.vSMC), T cells (TC), natural killer cells (NK), B cells and plasma cells (BC.PLASMA), macrophages and dendritic cells (MAC.DC), and monocytes (MONO) (Fig 2B and S1 Table). During subclustering analysis, we removed 113 endothelial cell and pericyte doublets as well as 26 epithelial and T cell doublet.

**Fig 2.**
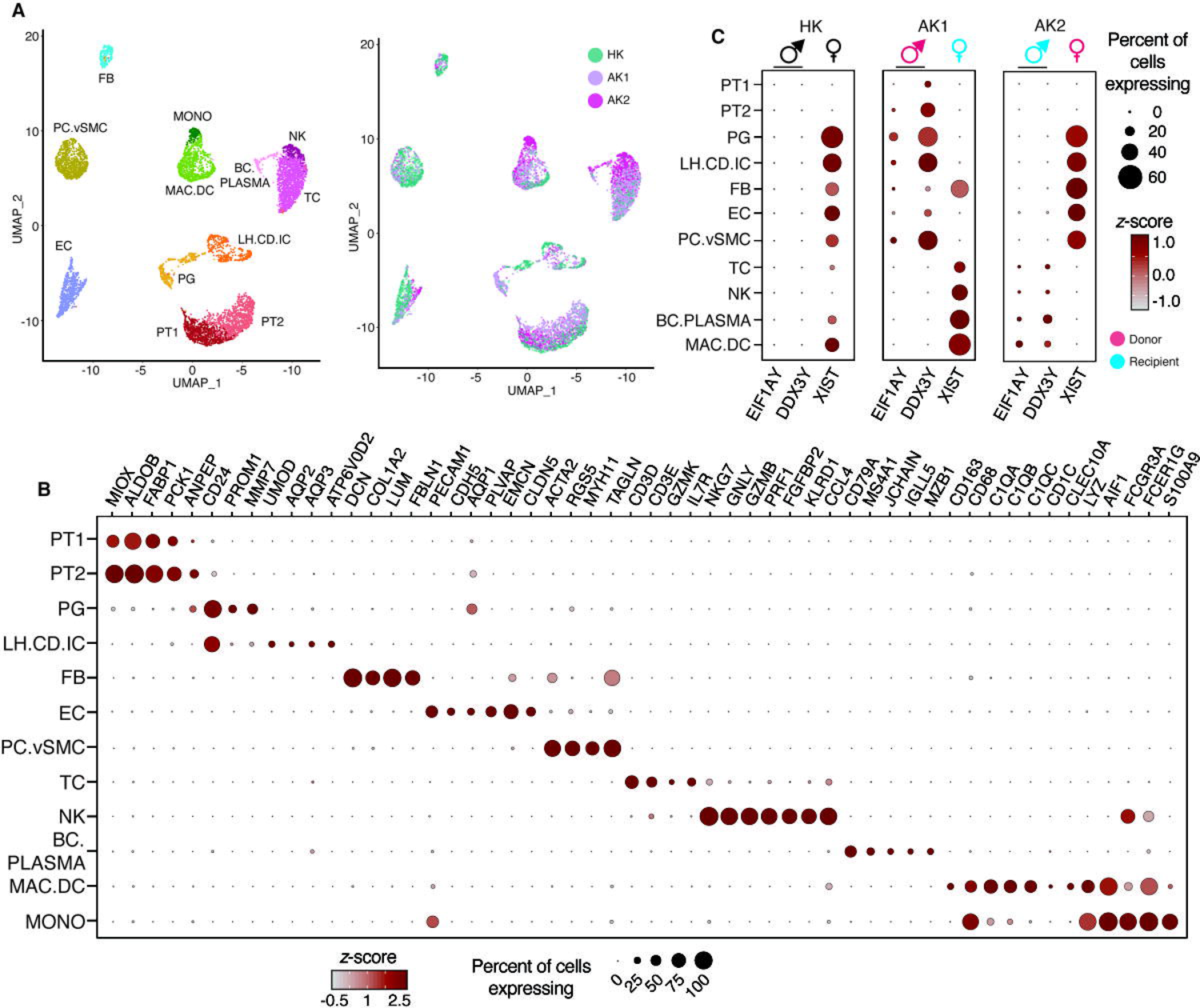
scRNA-seq applied to kidney allografts differentiated cell types and resolved recipient and donor origin based on sex-specific gene expression. (A). Uniform manifold approximation and projection (UMAP)-based visualization of 7217 individual cells obtained from the three kidney biopsy tissues. Left Panel: UMAP-based visualization in which different clusters represent different cell types. PT1 and PT2, proximal tubular cells 1 and 2; PG, progenitor cells; LH.CD.IC, cluster of loop of Henle, collecting duct, and intercalated cells; FB, fibroblasts; EC, endothelial cells; PC.vSMC, cluster of pericytes and vascular smooth muscle cells; TC, T lymphocytes; NK, natural killer cells; BC.PLASMA, cluster of B lymphocytes and plasma cells; MAC.DC, cluster of macrophages and dendritic cells; and MONO, monocytes. Right Panel: UMAP-based visualization of the same cell clusters shown in the left panel in which the cells are colored by the samples. HK-healthy kidney biopsy tissue, AK1 and AK2-allograft kidney biopsy tissues. (B). Dot-plot showing expression of known lineage markers. The size of the dot represents the proportion of cells within each cluster expressing the marker. The intensity of the color represents the standard score for each marker across different cell clusters. (C). Dot-plot showing the annotation of donor/recipient origin of cells in each sample based on female (*XIST*) and male (*EIF1AY*, *DDX3Y*)-specific gene expression patterns. HK is a female donor kidney; AK1 is a female recipient of a kidney from a male donor; AK2 is a male recipient of a kidney from a female donor. *XIST* is a female-specifically expressed gene. *XIST* gene transcription produces X-inactive specific transcript (Xist) RNA, a non-coding RNA, which is a major effector of the X chromosome inactivation. The Xist RNA is expressed only on the inactive chromosome and not on the active chromosome. Males (XY), who have only one X chromosome that is active, do not express it. Females (XX), who have one active and one inactive X chromosome, express it. In HK biopsy (female kidney), all the cells in the kidney express *XIST* and none express the Y chromosome markers. In AK1 biopsy (male donor and female recipient), all the kidney parenchymal cells express Y chromosome markers whereas all the recipient-derived immune infiltrating cells express *XIST*. In AK2 biopsy (female donor and male recipient), all the kidney parenchymal cells express XIST whereas all the recipient-derived immune infiltrating cells express the Y chromosome markers.

### Sex differences between kidney recipient and donor reveal migratory graft-infiltrating cells

AK1 was a female recipient of a kidney from a male donor and AK2 was a male recipient of a kidney from a female donor. We took advantage of the sex mismatch between kidney recipients and their donors and monitored the expression of male-specific Y chromosome-encoded *EIF1AY* and *DDX3Y* and female-specific *XIST*, involved in female X chromosome inactivation, to determine recipient or donor origin of the cells in the allograft [16]. We first separated the cell clusters by individual samples and then assigned each cluster to either female or male origin based on the expression pattern of *XIST*, *EIF1AY*, and *DDX3Y* genes. Overall, the frequency for capture of sex-specific transcripts was higher for female cells as XIST was more abundant in expression compared to *EIF1AY* and *DDX3Y*. HK biopsy was obtained from a female kidney donor and as expected, the cell clusters expressed XIST and were devoid for expression of *EIF1AY* and *DDX3Y*, indicating their female origin (Fig 1C). The assignment of the sex of cells was further improved by also including Y-chromosome autosomal-region-expressed *RPS4Y1* (S3 Fig).

In accord with AK1 being a female recipient of male kidney, all graft infiltrating immune cell types TC, NK, BC.PLASMA, and MAC.DC were of female origin and matched the sex of the allograft recipient. In accord with AK2 being a male recipient of female kidney, the graft infiltrating immune cell types were of male origin matching the sex of the recipient. In contrast to the infiltrating cells, the AK1 kidney parenchymal cells were male and matched the sex of the organ donor, while AK2 kidney parenchymal cells were female but also matched the donor. Importantly, XIST RNA expression was not universal across all female parenchymal cells as exemplified by absence in PT1 and PT2 in AK2 and HK female kidneys. The absence of *XIST* from female PT cells is a biological phenomenon rather than due to technical issues.

Unexpectedly, most of the FBs identified in male kidney AK1 were of female recipient origin (like the infiltrated immune cells), indicating that these FBs are migratory in nature and their presence in the allograft is by infiltration of cells from the recipient (S2 Table). The AK1 kidney biopsy had a Banff chronic lesion score of 2 for interstitial fibrosis (ci score), defined by interstitial fibrosis involving 26-50% of cortical area. This finding is particularly striking considering that FBs in AK2, where the biopsy had a Banff ci score of 0, defined by interstitial fibrosis involving ≤5% of cortical area, largely matched the donor (like the kidney parenchymal cells) and not the recipient.

### Comparative analysis of fibroblast-specific gene expression in healthy and allograft kidneys

To further resolve FB subtypes, we performed FB subclustering analysis and identified FB1 to FB4 (Fig 3A, top), all of which expressed the canonical fibroblast marker DCN (Fig 3B). The subtypes distinguish disease (FB1 and FB4) and healthy samples (FB2 and FB3) (Fig 3A, bottom). FB1 was comprised of all sex-mismatched (recipient-derived) ‘migratory FBs (migrating from the recipient/host to the donor/kidney allograft)’ (Fig 4A-C) whereas FB4 was comprised of sex-matched (donor-derived) ‘transplant-organ-originating’ fibroblasts of AK1 and AK2 biopsies. FB3 and FB4 were defined by expression of *GGT5* and *EMILIN1*, markers recently reported in healthy kidney biopsy to represent interstitial fibroblasts [29]. In addition, these FBs expressed *ACTA2*, indicative of myofibroblast (mFB)-like characteristics [29]. FB4 uniquely expressed *POSTN,* another mFB marker recently defined in a cell subpopulation in human kidneys expressing the most ECM genes [30]. Furthermore, FB4 cells uniquely expressed *TNC, COL4A1, COL18A1*, and *TGM2*, genes involved in beta-1 integrin cell surface interactions and cell adhesion to the extracellular matrix and included cellular stress response genes *JUN* and *FOSB* frequently observed in scRNA-seq due to stress of mechanical cell dissociation [31].

**Fig 3.**
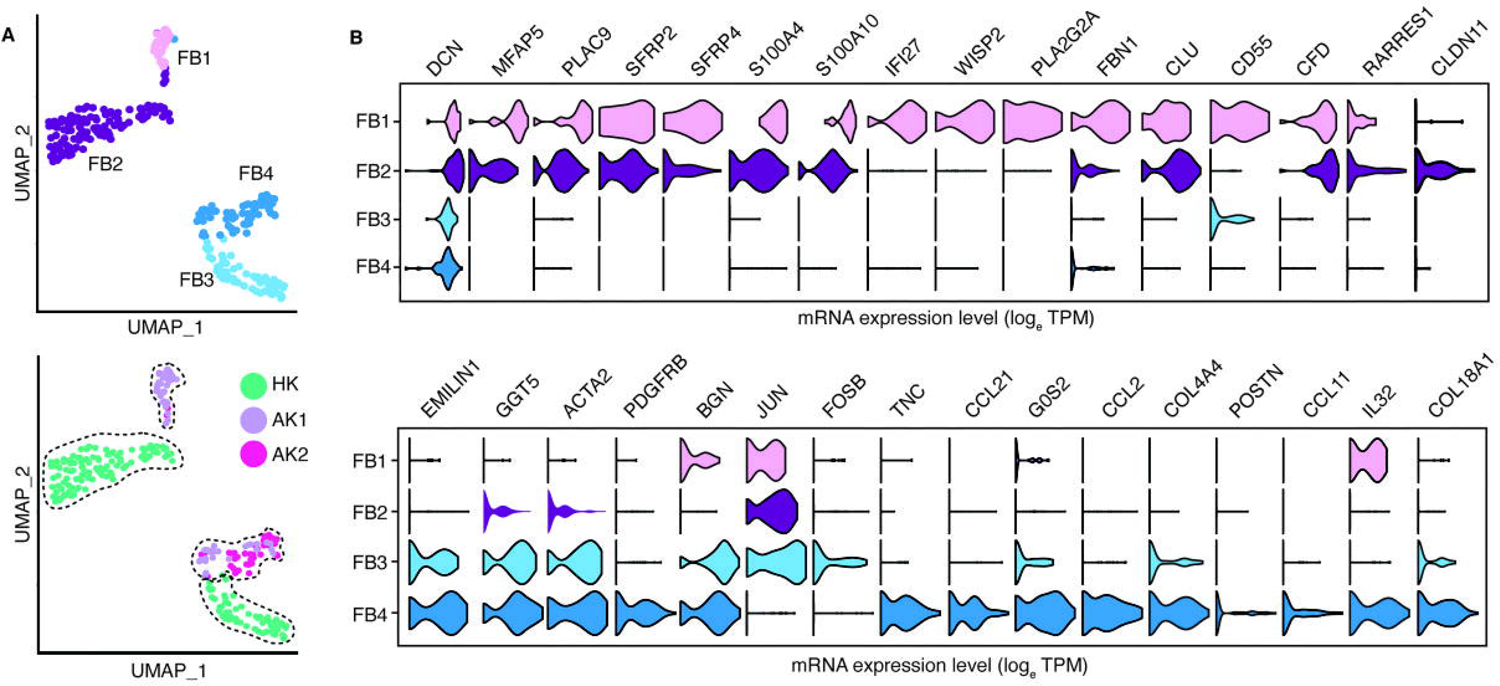
Sub-clustering of fibroblasts revealed four distinct subtypes. (A). UMAP-based visualization of subpopulations of fibroblasts. In the top panel the fibroblasts are colored by different sub-populations. In the bottom panel the cells are colored by the biopsies HK, AK1 and AK2. (B). Violin plots depict the expression of the lineage gene markers.

**Fig 4.**
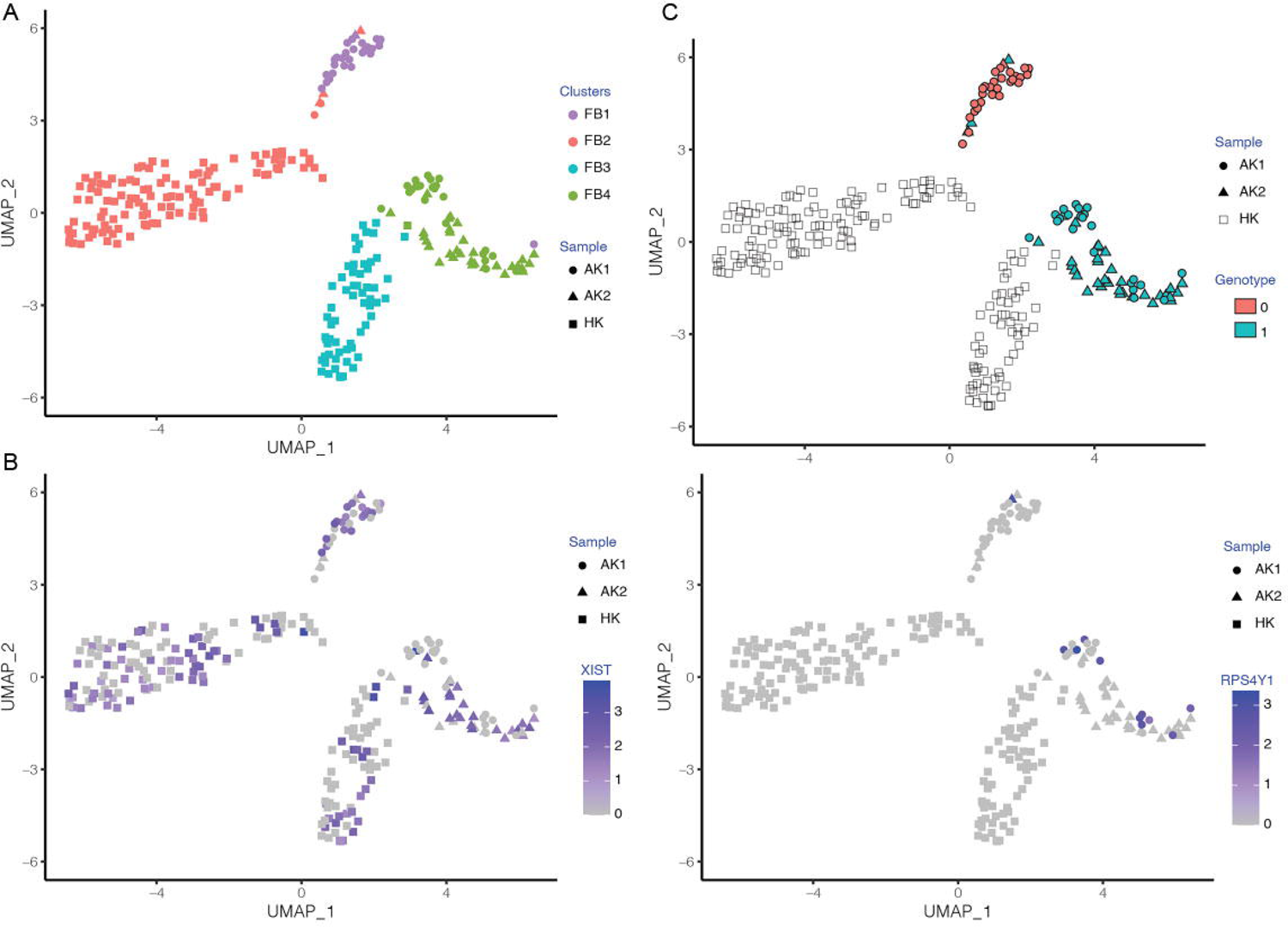
Visualization of fibroblast subpopulations clustered based on donor/recipient sex and clustered based on their genotype. (A). Fibroblast subpopulations. (B). Fibroblast subpopulations visualized based on their expression of female-specific gene XIST. (C). Fibroblast subpopulations visualized based on their expression of male-specific Y chromosome autosomal gene RPS4Y. (D). Fibroblast subpopulations visualized based on their genotype clusters

FB1 and FB2 expressed the secretory factors *CFD, SFRP2* and *SFRP4* in addition to genes implicated in promoting cell migration such as *MFAP5, S100A4* and *S100A10* [32]. The infiltrating FB1 cells uniquely expressed *FBN1*, *IFI27, WISP2* and *PLA2G2A*. The migratory FB1 expression signature was devoid of lymphoid or myeloid cell lineage markers suggesting that these cells are either not derived from hematopoietic lineage or their expression is lost in the target (donor) kidney. The absence of markers of other cell lineages also suggests these are not rare cell doublets (Fig 5). In addition, using an R package ‘DoubletFinder’ [33]—a computational doublet detection tool, we confirmed that there were no cell doublets in this migratory FBs. The migratory FB1 cells were comprised predominantly of female recipient-derived FB from AKI biopsy; however, we also detected 1 male recipient-derived migratory FB in the AK2 biopsy (Fig 4B-C). Clustering of the FB cells based on their genotype matched their clustering based on donor/recipient sex and confirmed the presence of recipient-derived migratory fibroblasts (Fig 4D). As expected, our FB cells mapped with a high prediction score to fibroblasts in the kidney reference dataset generated in the HuBMAP and KPMP projects (S4 Fig). Also, the expression of our FB lineage markers as well as the uniquely expressed genes of our migratory FB cells were largely restricted in the kidney reference dataset to FB cells (S5 Fig). Importantly, they did not map to any immune cells in the reference dataset, confirming the authenticity of our FB annotation. Both AK1 and AK2 biopsies contained exclusively kidney cortex and were not contaminated by kidney capsule or other extra-kidney tissues.

**Fig 5.**
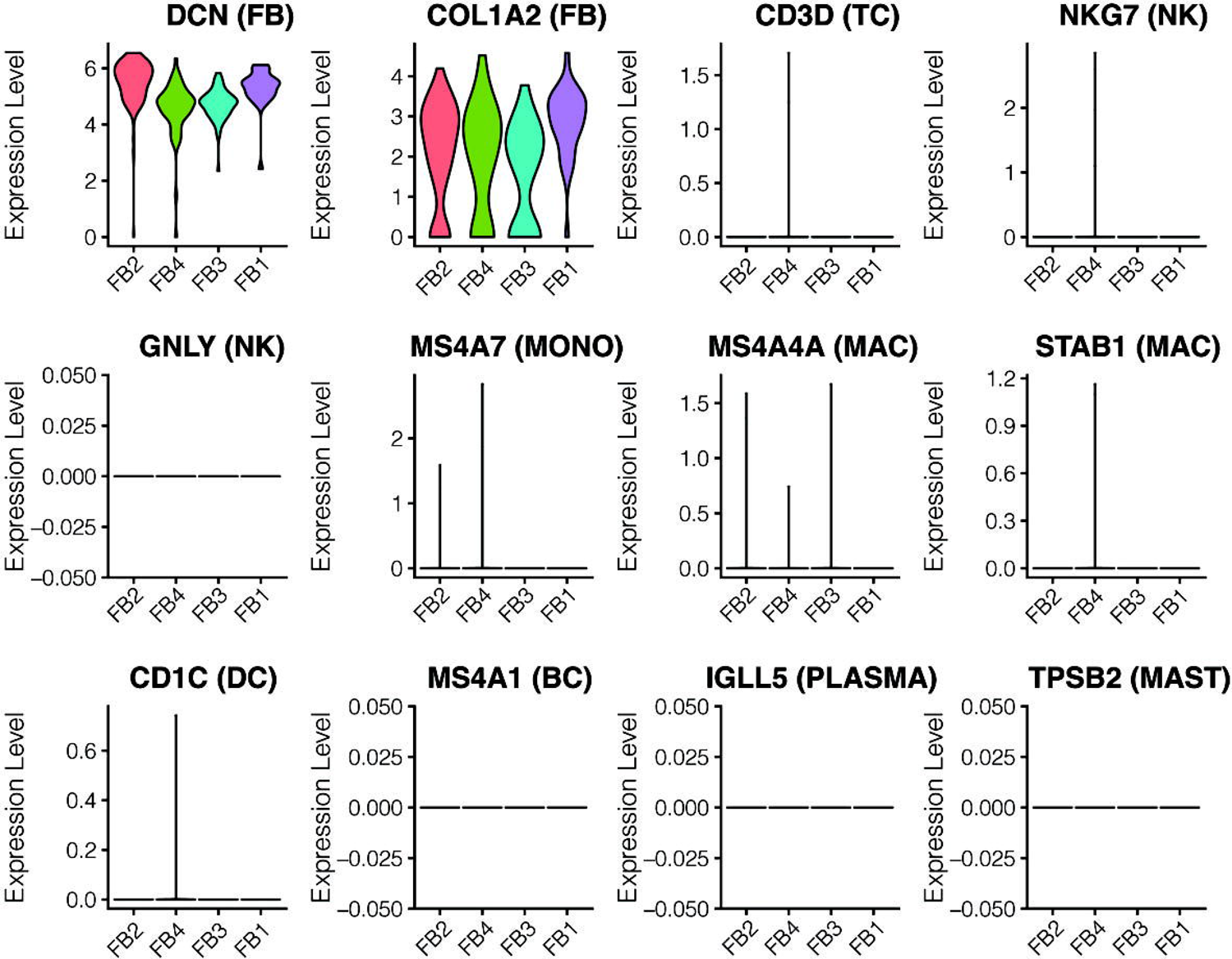
Expression of cell lineage markers in the fibroblast subpopulations Figure depicts the violin plots showing expression of cell lineage markers in the fibroblast subpopulations. No lymphoid or myeloid cell lineage markers were expressed in the fibroblasts.

### Validation of the four subpopulations of fibroblasts

Having defined four subpopulations of FBs including the migratory FB1, we compared our fibroblast subtype expression pattern with those of recently published kidney scRNA-seq studies (Fig 6). Subpopulation F3 and F4 expressing GGT5 and EMILIN1 (Fig 6A), were similar to kidney FBs (Fig 6B) observed in the study by Wu *et al.* [29] comprising a kidney allograft biopsy with histologic diagnosis of acute T-cell–mediated rejection with plasma cells and acute C4d-negative ABMR and a healthy kidney tissue biopsy from a discarded human donor kidney. Kidney allograft tissue was processed as a single cell and healthy kidney tissue was processed as a single nucleus for RNA sequencing. In addition, our FB4 subtype expressed *PDGFRA*, *PDGFRB* and *POSTN* similar to the ‘high-ECM expressing’ transitioning FB to mFB (Fig 6C) reported by Kuppe *et al.* [30]. This study included human kidney tissue from patients with normal kidney function or chronic kidney disease (CKD) due to hypertensive nephrosclerosis undergoing partial nephrectomy because of kidney cancer. In a recent study by Valenzi *et al.* [34] of human lung tissue from healthy controls and from patients with systemic sclerosis-associated interstitial lung disease, two major FB subpopulations were identified, including an *MFAP5^hi^* population in the lung with systemic sclerosis-associated interstitial lung disease (Fig 6D). The expression profile of our migratory FB1 fibroblasts strikingly matches with this *MFAP5^hi^* FBs sharing expression of *MFAP5*, *PLA2G2A*, *SLPI*, *CD34*, and *THY1* genes (Fig 6A and 6D). We next compared the expression profile of FB1 to FB4 with FBs of skin we reported previously [35]. Skin biopsies originated from patients with atopic dermatitis and normal subjects. All four subpopulations FB1-FB4 in the kidney including the migratory FB1 subpopulation, were distinctly different from skin FBs (S6 Fig).

**Fig 6.**
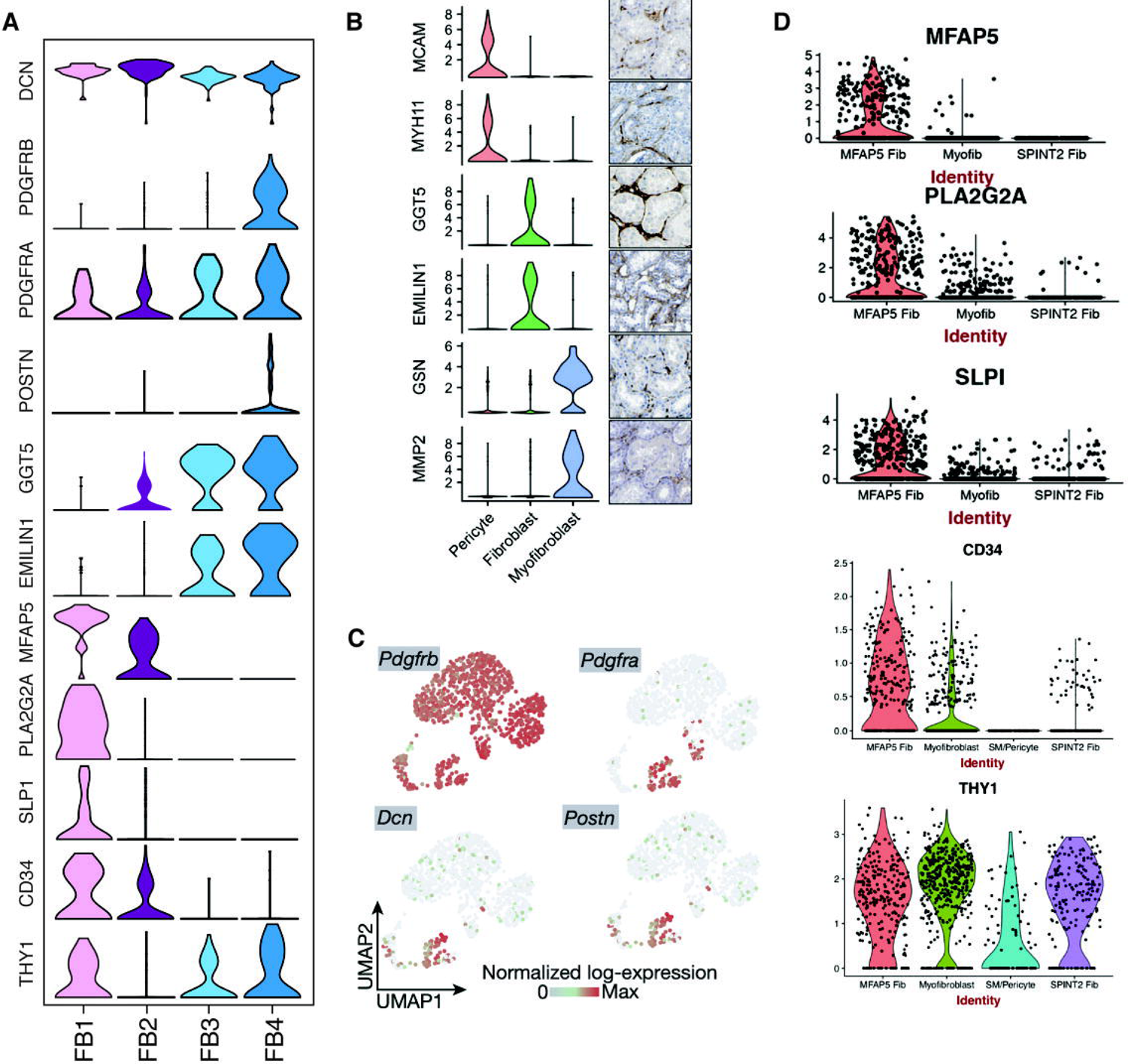
FB1-FB4 subpopulations compared with FBs described in recently published human scRNA-seq studies. (A). Violin plots depicting the expression of marker genes for FB1 to FB4 subpopulations. (B). FBs in a kidney allograft with rejection [29]. (C). FBs in kidneys with CKD due to hypertensive nephrosclerosis [30]. (D). FBs in lungs with systemic sclerosis-associated interstitial lung disease [34].

### Pericytes and vascular smooth muscle subtypes in the kidney

The vascular smooth muscle cell and pericyte cell types were further resolved into six subpopulations: vSMC1 to 4 and PC1 and 2 (Fig 7 and 8, S3 Table). The pericytes were characterized by *PDGFRA*-negative, *PDGFRB*-high, and *RGS5*-high signature (Fig 7B). vSMC1 and 2, were dominated by the HK biopsy, and showed abundant expression of stress-induced genes (relatively more in vSMC1 than vSMC2) such as *JUNB* and F*OSB* (Fig 7B). Interestingly, vSMC3 and 4, contributed by AK1 and AK2 biopsies, respectively, differentiated from the healthy vSMC1 and 2 by additional expression of cardiac muscle alpha actin *ACTC1* likely induced due to inflammation. vSMC3 was further characterized by induction of *NNMT,* considered as master metabolic regulator and contributor to secretion of cytokines and oncogenic extracellular matrix of cancer-associated fibroblasts [36]. The expression of *NNMT* also in PTC cells was unique to the AK1 allograft with high interstitial fibrosis. The only additional difference between vSMC3 and vSMC4 was the expression of male specific *EIF1AY*. A higher level of stress response genes including *JUNB* and *FOSB* separated PC1 from PC2. In addition, PC2 cells expressed *THY1*, *S100A4,* and *CCL2* and exclusively originated from HK cells.

**Fig 7.**
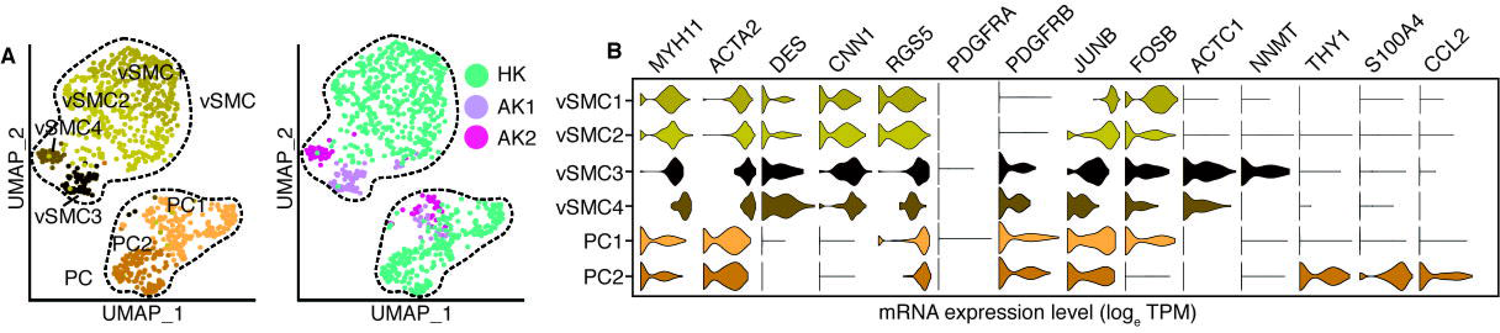
Sub-clustering of pericytes and vascular smooth cells. (A). UMAP-based visualization of subpopulations of pericytes and vascular smooth muscle cells. In the left panels the pericytes and vascular smooth muscle cells are colored by different sub-populations. In the right panel the cells are colored by the biopsies HK, AK1 and AK2. (B). Violin plots of the expression of lineage gene markers of pericytes and vascular smooth muscle cells.

**Fig 8.**
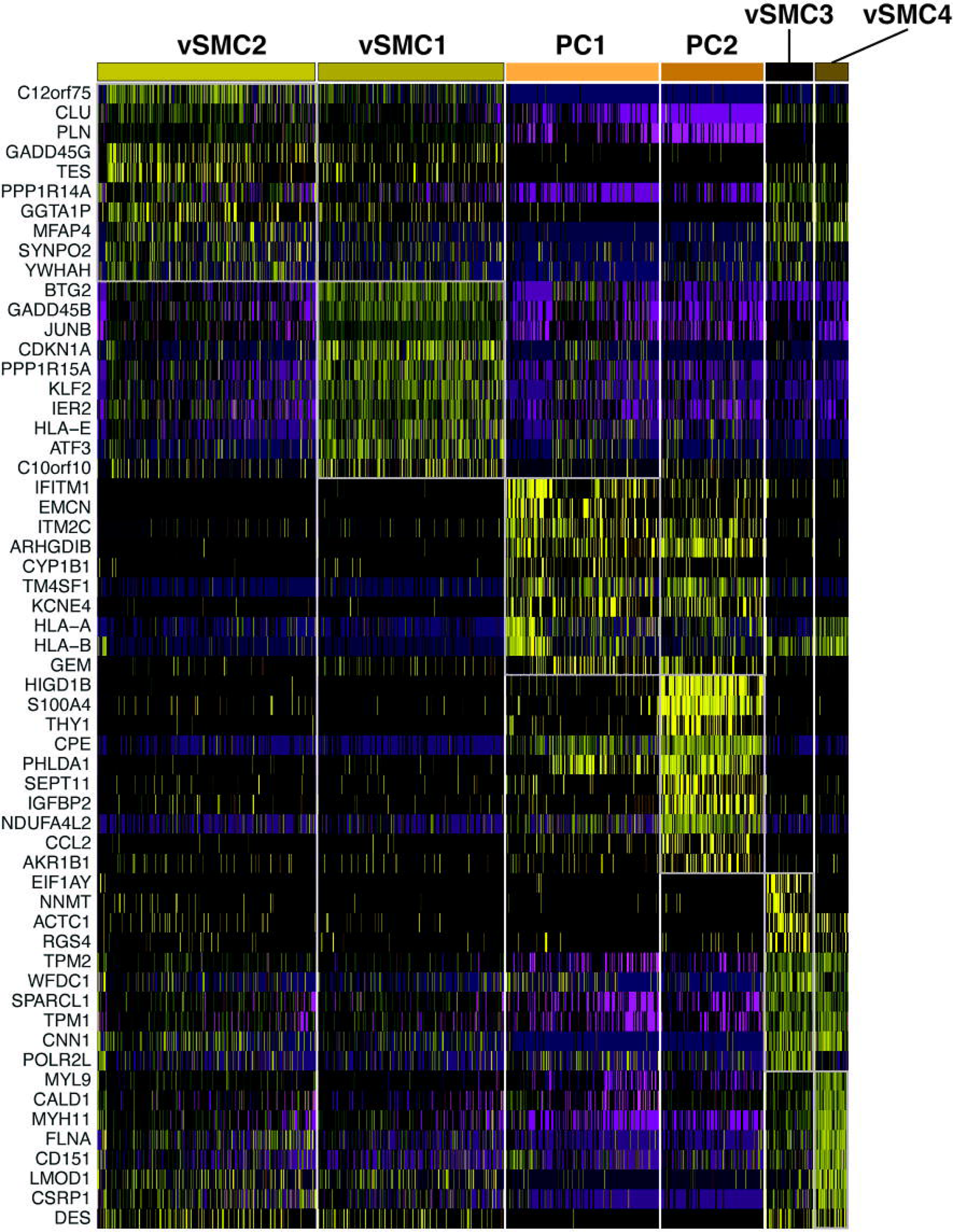
Differential gene expression in pericytes and vascular smooth muscle cells Figure depicts heatmap of differentially expressed genes in pericytes and vascular smooth muscle cells

### Tubular progenitor cell populations are increased in allograft kidneys

Reclustering analysis of the epithelial cell types PT1, PT2, PG, and LH.CD.IC yielded 6 subpopulations (Fig 9A-B). A single proximal tubular (PT) cell cluster was defined by expression of *MIOX, ANPEP,* and *SLC13A1* genes (Fig 9C). Tubular progenitor (PG) cells expressing *PROM1, CD24,* and *VIM*, however, separated into a major (PG1) and a minor (PG2) population. Fibrotic AK1 biopsy contributed the most to PGs (PG1 79.3% and PG2 55%) (Fig 9D). Heterogeneity in the proportion of kidney parenchymal cells among the three samples could be due sampling variability and should be interpreted with caution. PGs lack brush borders, are scattered throughout proximal tubules in the normal kidney and become more numerous and participate in tubular regeneration after acute tubular injury [37, 38]. We note that PGs also expressed *CDH2* (N-cadherin), a known marker for epithelial-mesenchymal transition (EMT) [39]. The co-expression of proximal tubular cell marker genes in PG1 suggests a more differentiated state compared to PG2. Furthermore, PG1 and PG2 are characterized by expression of injury markers *HAVCR1, LCN2,* and *MYC,* which are absent in PT (Fig 10).

**Fig 9.**
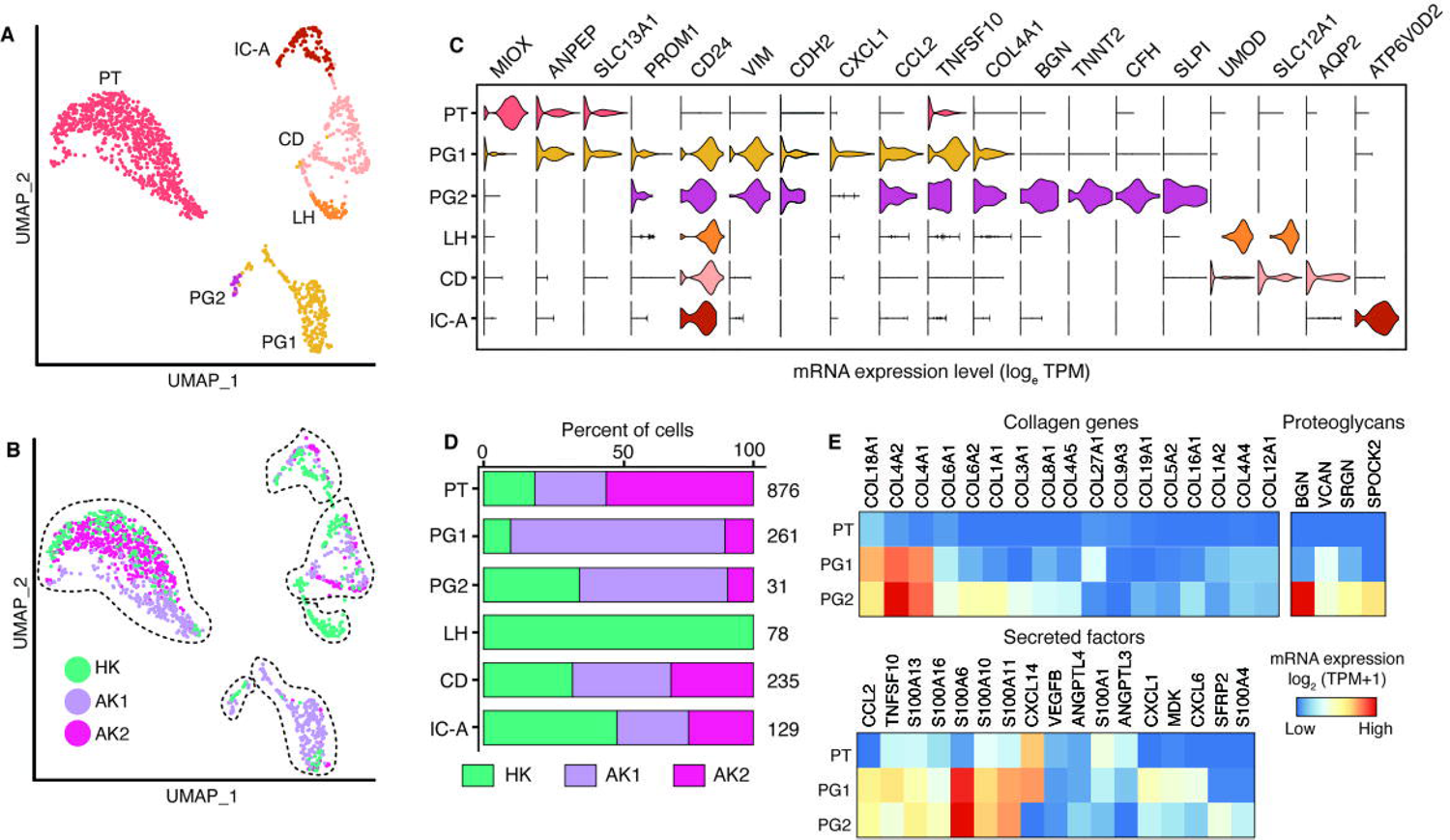
Epithelial cell sub-clustering revealed collagen-producing tubular progenitor cells. (A). UMAP-based visualization of epithelial cells colored by different cell types. PT, proximal tubular cells; IC-A, intercalated cells type A; CD, collecting duct cells; LH, loop of Henle cells; PG, progenitor cells. (B). UMAP-based visualization of epithelial cells colored by the biopsies HK, AK1 and AK2. (C). Violin plot showing expression of the lineage gene markers. (D). Stacked bar plots show the proportion of epithelial cells in each sample. The number on the right is the total number of cells. (E). Heatmap showing top expressed genes belonging to categories of matrisome groups.

**Fig 10.**
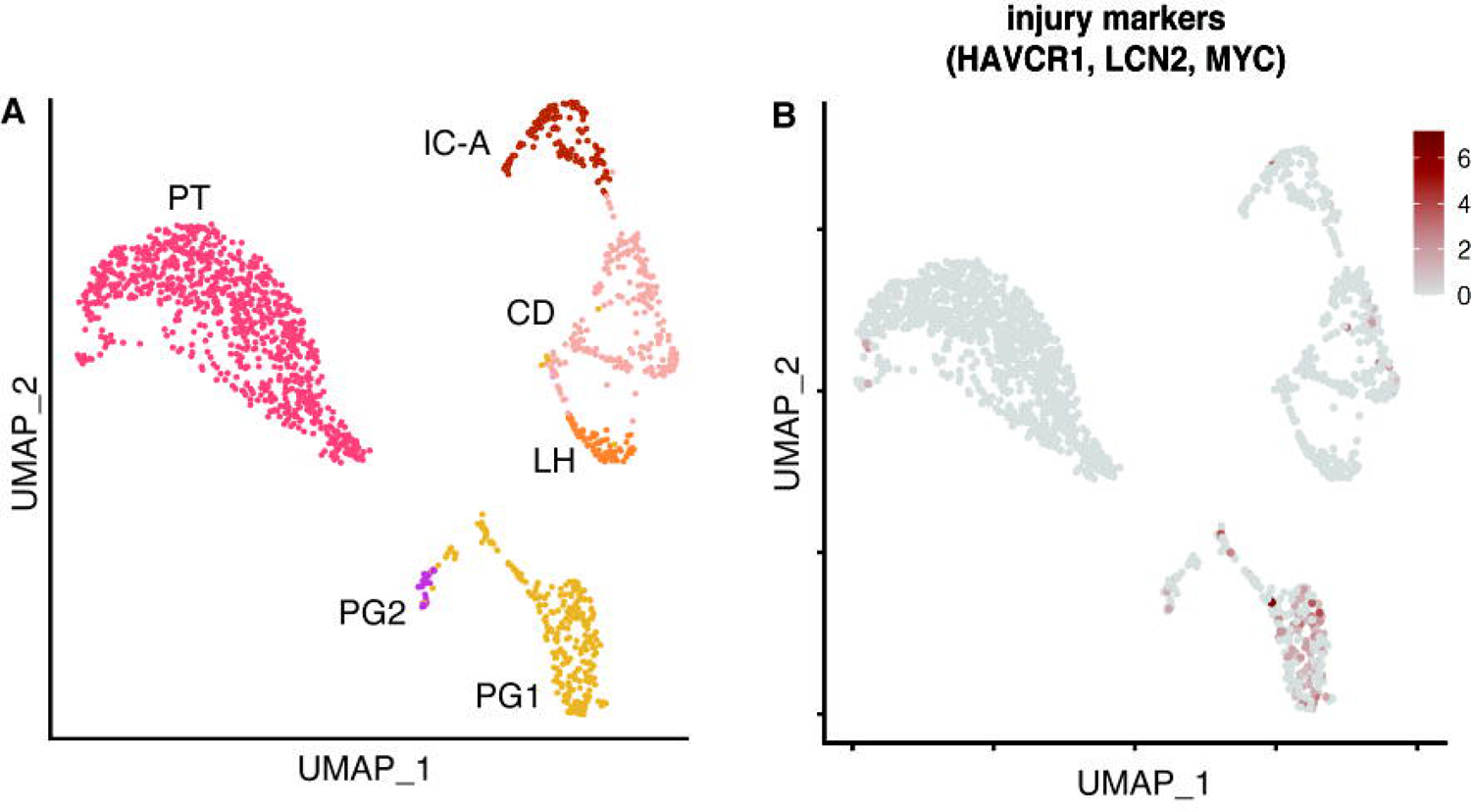
Analysis of kidney epithelial cell subpopulations Figure depicts UMAP-based visualization of epithelial cell groups. (A). UMAP-based visualization of epithelial cells. (B). Feature plot showing co-expression of kidney injury markers *HAVCR1, LCN2,* and *MYC*.

The key histological feature of AK1 kidney was interstitial fibrosis. To examine whether PG abundance and gene expression contributed to the fibrosis, we performed matrisome (the ensemble of extracellular matrix [ECM] and ECM-associated proteins) enrichment analysis [40, 41] of the PT and PG subtypes (Fig 9E) and also compared it to those of FB cells from our study (Fig 11). Most of the expressed collagen genes including *COL4A1, COL4A2,* and *COL1A1,* were selectively enriched in PGs and not or minimally expressed in PT cells. Similarly, the second category of matrisome genes ‘proteoglycans’ also showed enrichment in PGs and not in PT. In addition, secreted factors including the S100A family and cytokines such as *CCL2, CXCL1,* and *CXCL6* were abundant in PGs. The other epithelial cells of the kidney such as LH, CD, and IC-A were identified by unique expression of marker genes *UMOD, AQP2*, and *ATP6V0D2,* respectively.

**Fig 11.**
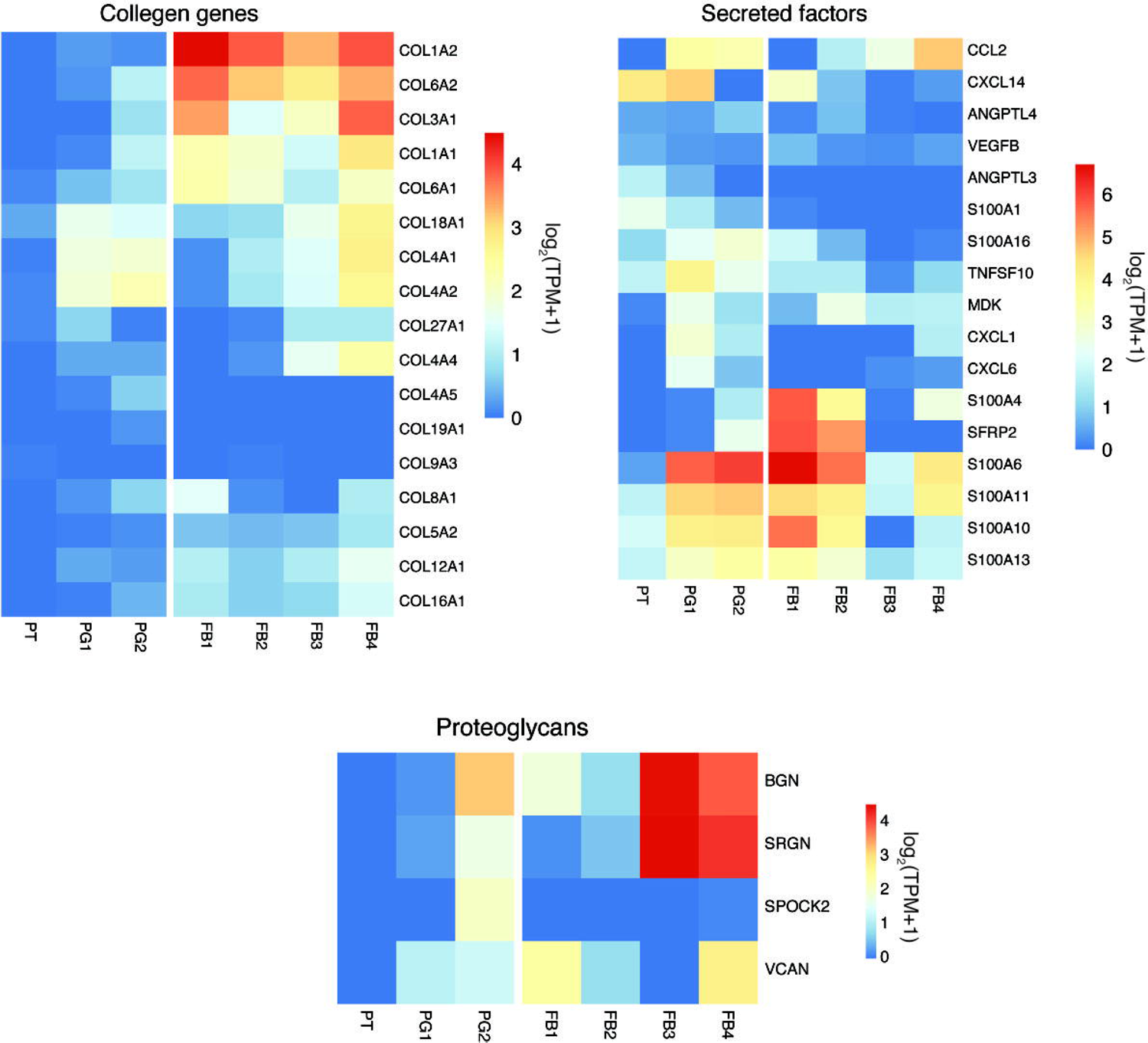
Top expressed genes of matrisome groups in fibroblasts, proximal tubular, and tubular progenitor cells Figure depicts heatmap of the top expressed matrisome genes in the proximal tubular cells, tubular progenitor cells, and fibroblasts.

### Endothelial cell types in disease biopsies show a chemoattractant cytokine signature

Further investigation of endothelial cell types revealed four distinct subtypes; peritubular capillaries (PTC1-3) and descending vasa recta (DVR), primarily based on the distinct expression for *PLVAP, AQP1,* and *SLC14A* (Fig 12A, B and S3 Table). Glomerular endothelial cells did not separate out by this analysis although some ECs co-expressed glomerular endothelial markers. All EC subclusters shared expression of canonical markers such as *PECAM1* and *CDH5.* The PTC2 subpopulation, mostly composed of AK2 cells, was characterized by the unique expression of the structurally and functionally related cytokines *CXCL9, CXCL10*, and *CXCL11* (Fig 12A-C). These cytokines act as chemoattractants during inflammation through binding to the receptor *CXCR3* mostly expressed by activated T cells [42]. AK1 and AK2 PTCs showed a >6-fold upregulation of the cell-surface-glycoprotein-encoding *SELE* gene compared to HK (*SELE* TPM, HK: 0.6, AK1: 2.6, AK2: 4.1). Under inflammatory conditions, endothelial cells induce expression of *SELE* in order to facilitate trans-endothelial passage of leukocytes through adhesion to the vascular lining [43]. The PTC3 subpopulation showed higher expression of *JUN, FOS* and *JUNB* presumably induced by mechanical stress of sample processing (Fig 13).

**Fig 12.**
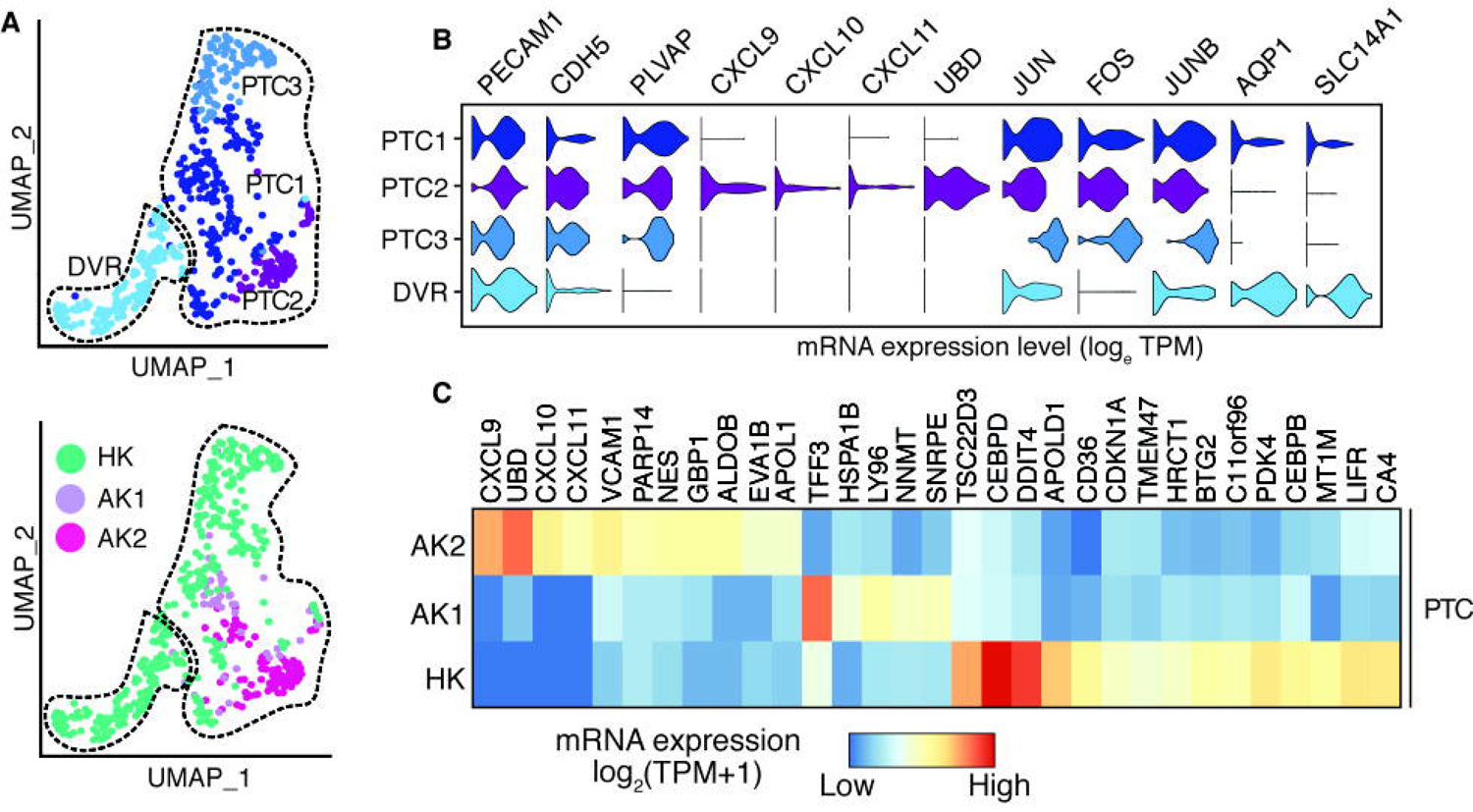
Sub-clustering of endothelial cells. (A). UMAP-based visualization of subpopulations of endothelial cells. In the left panel the endothelial cells, are colored by different sub-populations. PTC, peritubular capillaries; DVR, descending vasa recta. In the right panel the cells are colored by the biopsies HK, AK1 and AK2. (B). Violin plots of the expression of lineage markers of endothelial cells. (C). Heatmap shows the differentially expressed genes among the three biopsies in PTC cells (PTC1, PTC2 and PTC3 combined).

**Fig 13.**
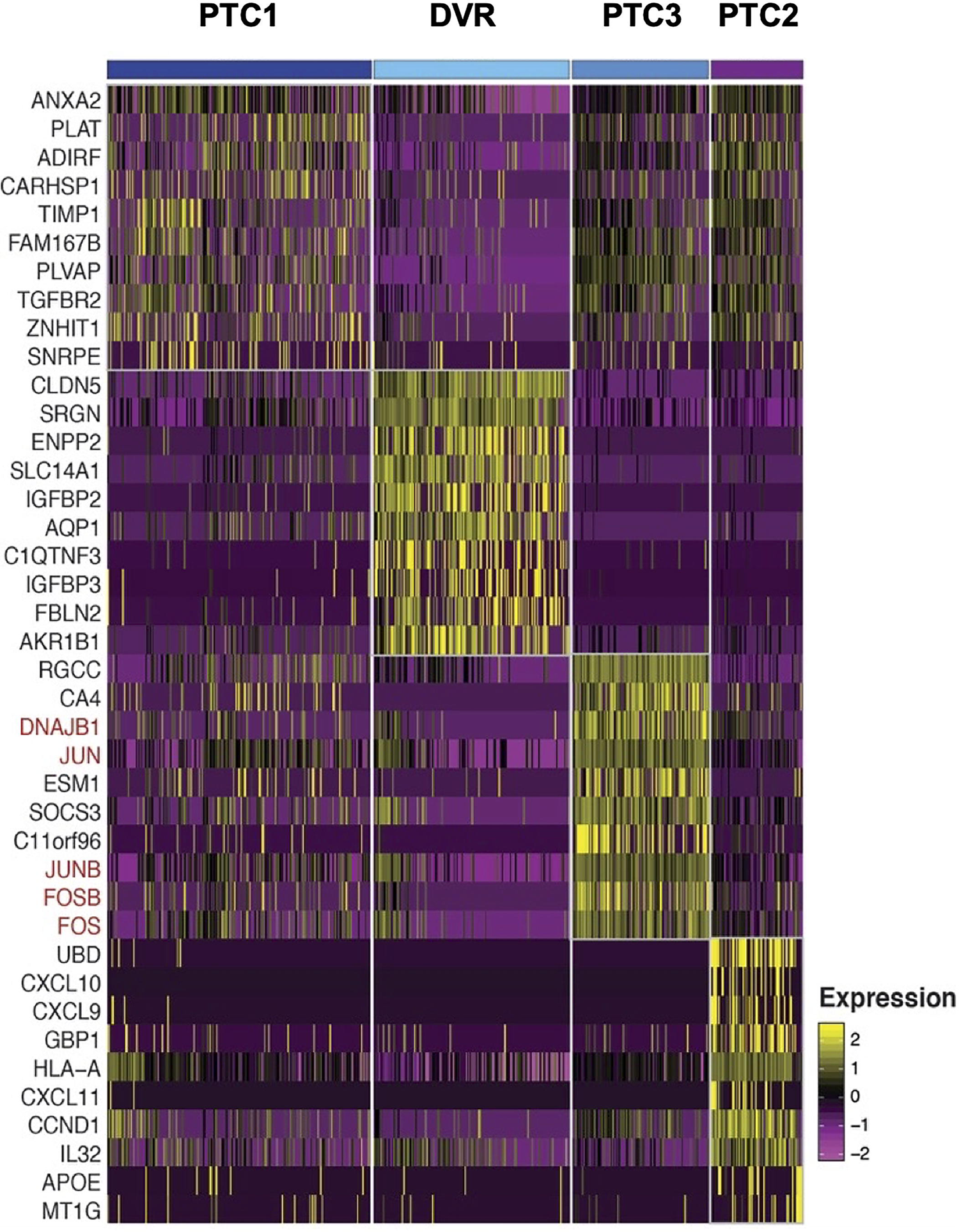
Differential gene expression in endothelial cell subtypes Figure depicts heatmap of the differentially expressed genes in the endothelial cell subtypes.

### Immune cell heterogeneity in the healthy and allograft kidneys

Subclustering analysis of the immune cell populations identified TC, cytotoxic TC, NK, MONO, MAC, DC, BC, PLASMA, and MAST (mast cells) cell types (Fig 14A-C). The transplant AK1 and AK2 samples showed higher proportions of immune cell infiltrates (52% and 66%, respectively, vs. 16% in HK of all cells) (Fig 14D). The T cells in HK were dominated by granzyme-K-(GZMK-) producing CD8^+^ T cells, and a small subset of interferon-stimulated-gene-(ISG-) high CD4^+^ T cells with increased expression of *ISG15, MX1, RSAD2, IFIT1*, and *IFIT2* (Fig. 14E). ISG-high CD4^+^ T cells were also found in AK1 and AK2. Furthermore, AK2 showed a large subpopulation of GZMK-expressing CD8^+^ cytotoxic T cells, and a smaller subgroup of CD8^+^ cytotoxic T cells, defined by high expression of granzyme B (*GZMB*) and perforin (*PRF1*). We also identified a minor population of central memory T cells in AK2 characterized by expression of *CCR7, SELL*, and *TCF7*. MACs expressed *MS4A4A, STAB1* and *SEPP1* typically considered as gene signatures of “alternatively activated” M2 macrophages. The content of NK and MONO in AK1 (0.38% and 0.44%, respectively) compared to AK2 (9.8% and 7.8%, respectively) was remarkably reduced. Interestingly, while only one PLASMA cell was detected in AK1, they represented about 1.5% of total cells in the AK2 sample. *IRF1* was ≥10-fold more abundant in MACs of AK2 (TPM 3.1) compared to AK1 (TPM 0.3) and HK (TPM 0.1). *IRF1* was also upregulated in cytotoxic TCs of AK2 (TPM 3.0) compared to AK1 (TPM 1.7) and HK (TPM 1.4). Given their small number, the biological relevance of immune cell heterogeneity should be interpreted with caution.

**Fig 14.**
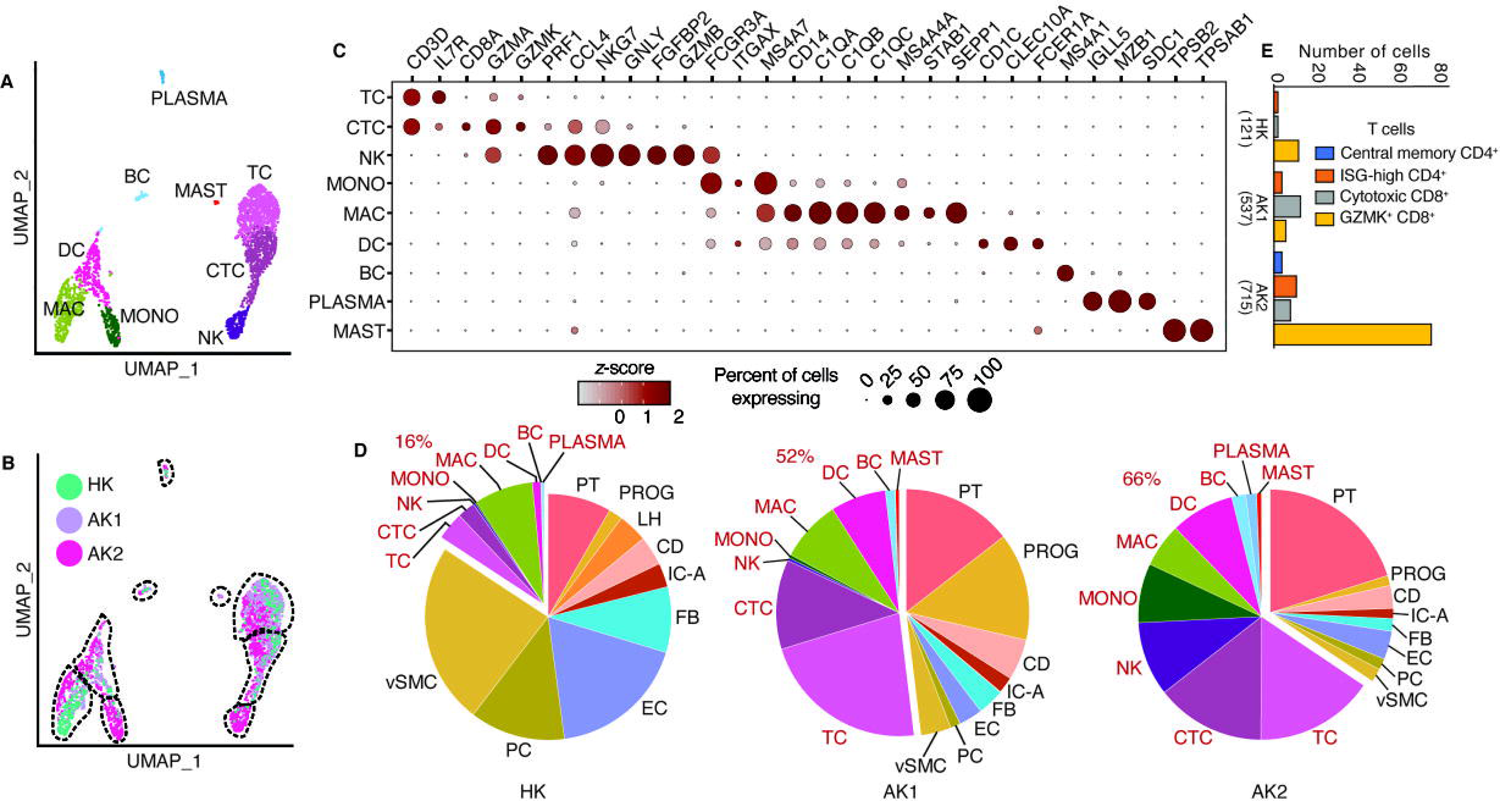
Immune cell heterogeneity in the healthy and allograft kidneys. (A). UMAP-based visualization of immune cells colored by different cell types. TC, T lymphocytes; CTC, cytotoxic T lymphocytes; NK, natural killer cells; MONO, monocytes; MAC, macrophages; DC, dendritic cells; BC, B lymphocytes; PLASMA, plasma cells. (B). UMAP-based visualization of immune cells colored by the biopsies HK, AK1 and AK2. (C). Dot-plot showing expression of known lineage gene markers. (D). Pie chart depicting the proportion of cell types in each biopsy sample. The immune cells are labelled in red. (E). Bar graphs showing the different types of T cells in HK, AK1, and AK2. Central memory CD4^+^ T cells: CCR7^+^SELL^+^TCF7^+^; ISG-high CD4^+^ T cells: CD4^+^ISG15^+^; Cytotoxic CD8^+^ T cells: CD8A^+^GZMB^+^.

## Discussion

The key findings of our single-cell transcriptome analysis include: (i) allograft kidney biopsy AK1, with ongoing tubulointerstitial fibrosis, contained more recipient-derived fibroblasts in contrast to the allograft kidney AK2 biopsy with no fibrosis and almost exclusively donor-derived FBs; (ii) allograft kidney AK1 biopsy also contained most proximal tubular progenitor cells that were enriched in the expression of ECM glycoproteins, collagens, and proteoglycans, and (iii) allograft kidney AK2 biopsy, eight months after successful treatment of antibody-mediated rejection as defined by clinical and histological criteria, contained endothelial cells expressing T cell chemoattractant cytokines. Determining the frequency of these observations is essential and will require scRNA-seq studies on larger cohorts of similar phenotype.

The discovery of migratory recipient-derived fibroblast subtypes in human allograft kidney biopsies is prompting for the investigation of molecular clues leading into recruitment of such migratory FBs to the kidney, the answer of which may hold immense therapeutic implications. In humans and in experimental kidney disease, macrophages can directly transdifferentiate into collagen-producing myofibroblasts [44]. However, the absence of immune cell or other cell lineage markers in our migratory FB1 subpopulation suggests that cells are either not derived from hematopoietic lineage or their expression is lost in the target (donor) kidney. The scRNA-seq study of human lung tissue from healthy controls and from patients with systemic sclerosis-associated interstitial lung disease identified two distinct major subpopulations of FB in the lungs [34]. One of the two subpopulation was called MFAP5^hi^ FB and was characterized by high expression of *MFAP5, CD34, THY1, SLPI*, and *PLA2G2A* in both healthy and diseased lungs [34]. Our migratory FB1 subpopulation showed high expression of *MFAP5, CD34, THY1, SLPI*, and *PLA2G2A*, qualifying as MFAP5^hi^ FBs found in lungs. This migratory FB1 subpopulation expressed fibronectin (*FBN1*), interferon-alpha-inducible protein 27 (*IFI27*), WNT1-inducible signaling pathway protein 2 (*WISP2*), and complement-decay-accelerating factor (*CD55*) uniquely. All these genes have been implicated in cancer cell migration and metastasis [45–48]. Increased *MFAP5* expression has been reported to stimulate cancer cell motility and invasion and predicted poor survival, and *MFAP5* in vivo silencing reduced tumor progression [32]. In addition to *MFAP5*, migratory FB1 identified in the AK1 biopsy were differentiated from the non-migratory FB4 subpopulation by *S100A4* expression in the migratory FBs. Interstitial fibroblasts appear after gastrulation because of EMT from secondary epithelium [49]. Calcium-binding protein *S100A4*—also called FSP1—likely plays an important role in facilitating and maintaining EMT phenotype [50]. *S100A4* is also expressed in some endosteal lining cells and marrow stromal cells and may be an EMT niche for osteogenic precursor cells, indifferent endosteum, fibroblasts, and marrow stromal cells [50]. Additionally, *S100A4* promotes metastasis and is associated with intestinal fibroblast migration in patients with Crohn’s disease [51, 52]. Our observation suggests that the migratory and tissue-invasive nature of the FBs may play a crucial role in fibrosis and complements the prior observation of mesenchymal cells of host origin in the vascular and interstitial compartments of kidney allografts undergoing chronic rejection [53]. The discovery of migratory FBs expressing unique genes associated with cell mobility and migration and may provide opportunities for targeted intervention to ameliorate interstitial fibrosis.

A recent scRNA-seq analysis of study of human kidney tissue collected from patients with normal kidney function or CKD due to hypertensive nephrosclerosis undergoing partial nephrectomy because of kidney cancer sought to identify the cell types that contribute to the production of extracellular matrix (ECM) in kidney fibrosis [30]. Among the four major cell types in the kidney—mesenchymal, epithelial, endothelial, and immune—mesenchymal cells exhibited the highest extracellular matrix expression in the kidney with minor contribution from epithelial cells. Within the mesenchymal cell type, 3 fibroblasts 2 myofibroblasts, 2 pericytes, and 1 vascular smooth muscle cells were identified. myofibroblasts (mFB) were defined as cells that expressed the most ECM genes and expressed the marker *POSTN.* Three main sources of mFB were identified—*NOTCH3^+^ RGS5^+^PDGFRA^-^* pericytes, *MEG3^+^ PDGFRA^+^* fibroblasts, and *COLEC11^+^CXCL12^+^* fibroblasts [30]. Using such a definition, our FB4 subpopulation qualified as mFB. In both AK1 and AK2 biopsies, FB4 subpopulation expressed markers that suggests contribution from all three sources of mFB. In our native kidney HK biopsy with no fibrosis, both FB2 and FB3 did not express *POSTN* and were likely not derived from pericytes.

If mFB may be defined as a cell population that express the most ECM genes and express the marker *POSTN*, then the source(s) of such cells in the kidney continues to remain controversial [54]. These cells were traditionally thought to arise from kidney resident interstitial FB—residual embryonic mesenchymal cells left over from organogenesis [55]. Data from more recent preclinical models suggests that they may originate from EMT [50], endothelial-to-mesenchymal transition [56], pericytes [57], and bone marrow-derived fibroblast precursor fibrocytes [58]. However, their relative contribution is not known and may depend on the nature of injury resulting in fibrosis [59]. Based on our findings, we speculate that kidney resident interstitial FB are the dominant cells that produce ECM in normal healthy kidney. In kidney allografts, in the context of alloimmune injury and toxicity due to drugs such as calcineurin inhibitors, there is additional contribution of pericyte-derived mFB to the pool of ECM-producing cells. As allograft fibrosis worsens, there is further recruitment of FBs to the pool from the recipient. The striking similarity between expression profile of our migratory FB1 and the *MFAP5^hi^* FB in the lungs suggests a common origin of these fibroblasts with potential role to affect multiple organs. Confirming our findings is essential and will require scRNA-seq studies on larger cohorts of similar phenotype. In this context, it will be interesting to study allografts with chronic active T-cell-mediated rejection or interstitial fibrosis and tubular atrophy not otherwise specified, and well-functioning allografts over the long term that never had acute rejection to assess whether recipient-derived migratory FBs are operational in those conditions as well.

The detection of *PROM1-(CD133-)* positive tubular progenitor (PG) cells expressing ECM glycoproteins, collagens, and proteoglycans in AK1 biopsy with interstitial fibrosis and tubular atrophy is a novel finding. While the contribution of EMT to kidney fibrosis is a subject of considerable controversy, recent evidence suggests that partial EMT is sufficient to induce the release of fibrogenic cytokines [60, 61]. The expression of *CDH2* (N-cadherin) in PGs and not any other tubular epithelial cell type, suggests PGs undergo partial EMT. The expression of injury markers seen in PGs suggest that chronic tubular injury likely induces EMT transition leading to production of extracellular matrix proteins and contribute to kidney fibrosis. Furthermore, although PGs express tubular cell markers, they could not be matched to previously established tubular cell subsets. PGs express several S100 proteins, a family of calcium-binding proteins involved in cell apoptosis, migration, proliferation, differentiation, energy metabolism, and inflammation. *PROM1^+^* cells are known to be distributed throughout the kidney and capable of expansion and limited self-renewal [62]. Identifying partial EMT in tubular progenitor cells rekindles the role of tubular cells in perpetuating fibrosis [63].

A subpopulation of endothelial cells in AK2 biopsy expressed mRNA for cytokines *CXCL9, CXCL10* and *CXCL11* while T cells expressed the cognate receptor *CXCR3*. Our finding of endothelium-derived T-cell chemoattractant cytokines provide a mechanistic basis for the presence of T cells in the AK2 biopsy. Donor endothelium derived *CXCL10* as initiator of alloresponse has been previously noted in a cardiac allograft model [64]. In kidney transplant recipients, urinary *CXCL10* levels increase during antibody-mediated rejection that is accompanied by microvascular inflammation [65]. However, to our knowledge, endothelial cells as a source of *CXCL9,10,11* has not been previously described in human kidney allografts. Such communication of endothelial cells through T cell chemoattractant, eight months after successful treatment of antibody-mediated rejection, is striking and highlights the role of donor-derived endothelium in perpetuating tissue injury in the presence of circulating antibodies directed against the donor HLA. Importantly, in this recipient, even though there was clinical and histological improvement, antibodies directed against the donor HLA persisted in the circulation.

Unlike donor and recipient FB chimerism, we did not find donor and recipient immune cell chimerism in the kidney allograft. Our findings of the absence of donor-derived T-lymphocytes in the kidney allograft agrees with the observation that donor T-lymphocyte number in the allograft decreased as a function of time, in a recent study that utilized whole exome sequencing of donor and recipient DNA, and scRNA-seq of the kidney allograft to identify the origin of each cell [66]. This study also found that recipient-derived macrophages dominate during rejection, and in a single patient who had a biopsy after treatment of rejection—33 months after transplantation, the proportion of donor-derived macrophages increased. In contradistinction, the lack of donor macrophages in our post-rejection treatment AK2 biopsy—performed at 84 months after transplantation—suggests the possibility that donor macrophages in the allograft also decreases as a function of time after transplantation.

A strength of our study is the rapid processing of freshly collected biopsies without cryopreservation, albeit on different days. We minimized the time for transfer of the samples from the ultrasound suite or the operating room, where the biopsies were performed, to the research laboratory for generating single-cell suspensions, and from the research laboratory to the genomics core laboratory for library preparation. We used the 10x Chromium platform (10x Genomics) with high cell-capture efficiency and permitting the use of human kidney allograft biopsies with limiting amount of tissue [6]. Earlier reports on scRNA-seq of healthy and diseased human kidney tissue, both native and allograft kidney, have included a combination of single-cell and single-nucleus sequencing [29], fresh and frozen specimens [67], multiple platforms to capture the single cells [29, 68] and analyzing the transcriptome after combining cells obtained from biopsies from tumor nephrectomies and discarded donor kidneys [69]. Another strength is the use of normal kidney tissue from a living kidney donor, instead of unaffected areas of tumor nephrectomies or kidneys rejected for clinical transplantation.

Our study has limitations. The number of study subjects was sparse, and we analyzed only three biopsies. Our findings need to be validated in more biopsies. Extrapolation from these results is challenging because of the heterogeneity in allograft pathology and tissue sampling depth by core needle biopsy. We did not study chronically active T-cell-mediated rejection or allografts with no acute rejection and stable function several years after transplantation. However, having implemented a standardized protocol for sample accusation, sample preparation, single-cell capture, RNA-seq, and data analysis, we believe that our results are robust and have provided rich mechanistic information. Another limitation is that not every cell type is captured by our whole cell dissociation protocol, in particular podocytes. Nevertheless, we were able to capture and analyze several nephron cell types; continued refinement of tissue processing techniques is expected to further improve the types of cells captured. Finally, there are inherent limitations to the scRNA-seq technique, for example, only a fraction of mRNAs present in cells are captured and converted to cDNA and the tissue dissociation process disrupts tissue architecture and loses relative spatial positioning information of cells.

## Conclusions

We have demonstrated the utility of scRNA-seq in interrogating intragraft events in kidney allografts. Our analysis has revealed unique cell types and cell states in kidney allograft biopsies and confirmed our hypothesis that applying scRNA-seq furthers precision transplantation medicine approaches by providing mechanistic insights and opportunities for drug target and pathway identification at hitherto unknown and unavailable resolution. With improvement in the technology, refinement in computational approaches, and decreasing operational costs, it is possible in the future to apply single-cell transcriptomics to complement conventional histopathology, in the clinic, for the idealized care of transplant recipients.

## Acknowledgements

We thank ThuTrang Du, Division Administrator, Division of Nephrology, Weill Cornell Medical College, for her help with the execution of this project. We thank A. Hurley and the Research Facilitation Office staff at Rockefeller University for their regulatory and administrative assistance.

## Supporting Information

S1 Fig. Histopathological characteristics of kidney allograft biopsy AK1

S2 Fig. Histopathological characteristics of kidney allograft biopsy AK2

S3 Fig. UMAP-based visualization of individual cells after including the Y chromosome gene

S4 Fig. Prediction scores for cell clusters when mapped to a kidney reference dataset

S5 Fig. Agnostic mapping of cells from our study to the kidney reference dataset

S6 Fig. Comparison of kidney fibroblasts with skin fibroblasts

S1 Table. Cell-type-specific gene expression

S2 Table. Cell-type-specific expression of the X chromosome marker gene XIST

S3 Table. Differential gene expression in subclustering analysis

