## Supplementary material for "Detection of infiltrating fibroblasts by single-cell transcriptomics in human kidney allografts": S1 fig

### S1 Fig. Histopathological characteristics of kidney allograft biopsy AK1

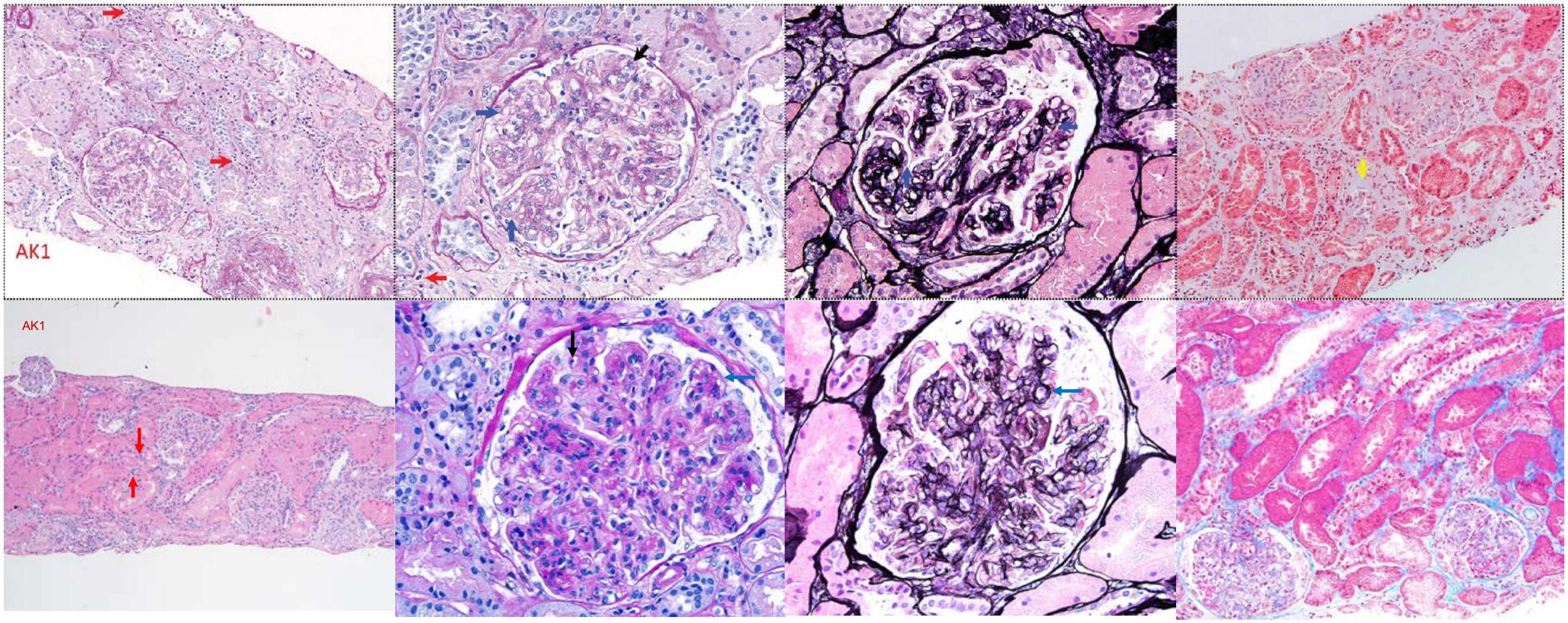


**Top:** Index biopsy used for scRNA-seq—done 42 months after kidney transplantation.
**Bottom:** Biopsy 16 months prior to the index biopsy—done 26 months after kidney transplantation.

Photomicrographs show periodic acid–Schiff stain at 200 (left) and 400 (center left) magnification, Jones methenamine silver stain at 400 (center right) and Masson's trichrome stain at 200 (right) magnification.

Biopsy (top) with severe transplant glomerulopathy(cg3), severe microvascular inflammation (g3, ptc3), and moderate tubulointerstitial fibrosis (ci2, ct2), was from a female patient with lupus nephritis of her native kidneys, 42 months after transplantation and 16 months after a prior biopsy (bottom) that showed severe transplant glomerulopathy (cg3), severe glomerulitis (g3), and no tubulointerstitial fibrosis (ci0, ct0).

Red arrow: peritubular capillary inflammation; Black arrow: glomerular inflammation; Blue arrow: duplication of glomerular capillary basement membrane; and Yellow arrow: interstitial fibrosis and tubular atrophy**.**
