## Supplementary material for "Detection of infiltrating fibroblasts by single-cell transcriptomics in human kidney allografts": S2 fig

**S2 Fig. Histopathological characteristics of kidney allograft biopsy AK2**

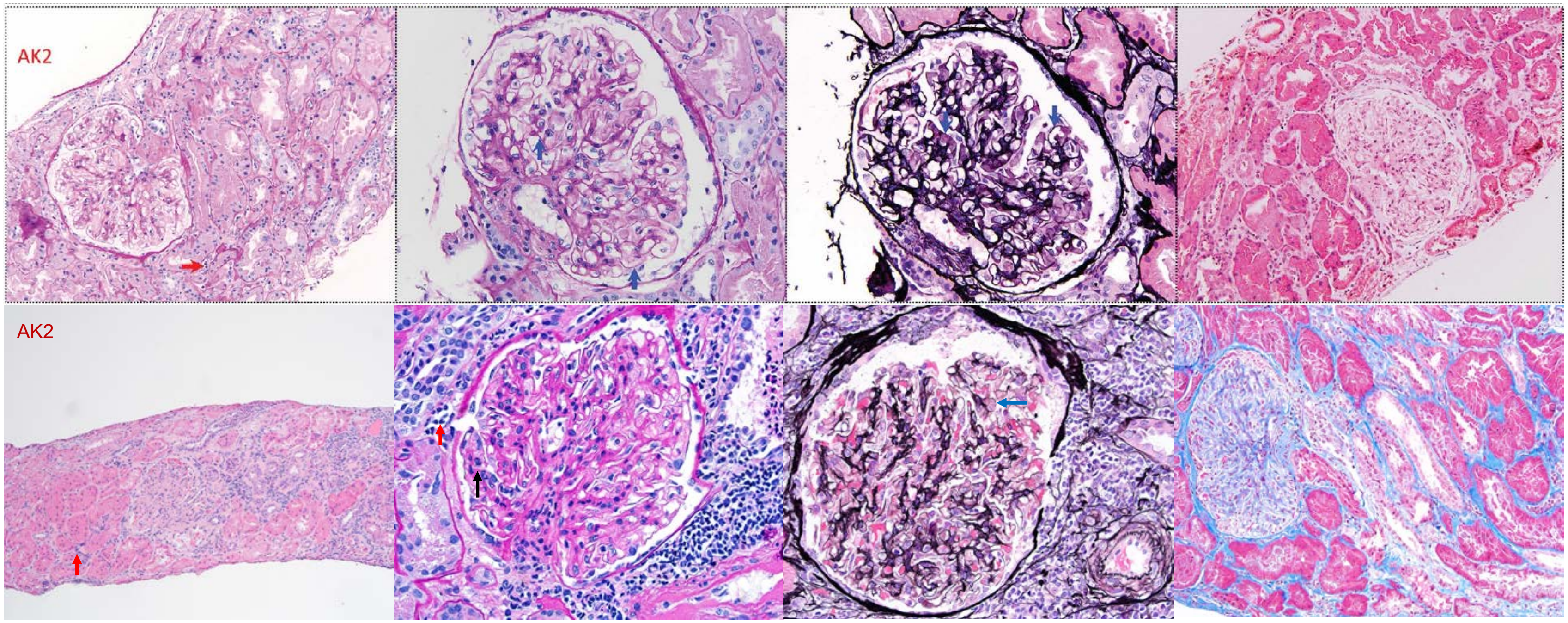


**Top:** Index biopsy used for scRNA-seq—done 84 months after kidney transplantation.
**Bottom:** Biopsy 8 months prior to the index biopsy—done 76 months after kidney transplantation.

Photomicrographs show periodic acid–Schiff stain at 200 (left) and 400 (center left) magnification, Jones methenamine silver stain at 400 (center right) and Masson's trichrome stain at 200 (right) magnification.

Biopsy (top) with minimal interstitial (i1, t0) and peritubular capillary inflammation (ptc1), was from a male patient with focal segmental glomerulosclerosis, 84 months after transplantation and 8 months after a prior biopsy (bottom) that showed active antibody mediated rejection (g2, ptc3, cg0, ci0, ct0)

Red arrow: peritubular capillary inflammation; Black arrow: glomerular inflammation; and Blue arrow: normal glomerular capillary basement membrane.
