## Supplementary material for "Detection of infiltrating fibroblasts by single-cell transcriptomics in human kidney allografts": S2 table

### S2 Table. Cell-type-specific expression of the X chromosome marker gene XIST

| Cell type | HK | Cell type | AK1 | Cell type | AK2 |
|---|---|---|---|---|---|
| PT | 0 | PT | 0.026925 | PT | 0.063701 |
| PG | 3.868948 | PG | 0.018473 | PG | 3.491369 |
| LH | 9.334275 | CD | 0.029711 | CD | 4.220906 |
| CD | 5.450415 | IC.A | 0 | IC.A | 3.703303 |
| IC.A | 4.310801 | FB1 | 3.530406 | FB4 | 4.782507 |
| FB2 | 2.440544 | FB4 | 0.675691 | AVR | 3.926241 |
| FB3 | 3.04712 | AVR | 0.290284 | vSMC | 3.18347 |
| AVR | 5.393421 | vSMC | 0 | PC | 3.794725 |
| DVR | 4.014057 | PC | 0 | TC.CTC | 0 |
| vSMC | 3.586586 | TC.CTC | 3.808262 | NK | 0 |
| PC | 4.797342 | MAC | 4.314509 | MONO | 0.017436 |
| TC.CTC | 2.325882 | DC | 3.854761 | MAC | 0 |
| MAC | 5.780216 | BC | 9.149131 | DC | 0.029137 |
| DC | 3.219575 | | | BC | 0 |
| | | | | PLASMA | 0 |
| | | | | MAST | 0 |

- Cell types with greater than 10 cells are shown
- Numbers represent TPM values
- AK1: Donor - male; Recipient - female
- AK2: Donor - female; Recipient - male

Immune cells

Absolute expression values in transcript per million (TPM) of XIST gene. XIST gene produces X-inactive specific transcript (Xist) RNA, a non-coding RNA that is a major effector of the X chromosome inactivation. The Xist RNA is expressed only on the inactive chromosome and not on the active chromosome. Males (XY), who have only one X chromosome that is active, do not express it. Females (XX), who have one active and one inactive X chromosome, express it. In healthy kidney HK (female kidney), all the cells in the kidney express XIST and none express the Y chromosome markers. In AK1 biopsy (male donor and female recipient), all the kidney parenchymal cells express Y chromosome markers whereas all the recipient-derived immune infiltrating cells express XIST. In AK2 biopsy (female donor and male recipient), all the kidney parenchymal cells express XIST whereas all the recipient-derived immune infiltrating cells express the Y chromosome markers. Thus, in the three biopsies, the X and Y chromosome markers are expressed as expected. Interestingly, the fibroblasts in AK1 biopsy (male donor and female recipient), expressed XIST, proving that these were recipient derived.
