## Supplementary material for "Detection of infiltrating fibroblasts by single-cell transcriptomics in human kidney allografts": S3 fig

**S3 Fig. UMAP-based visualization of individual cells after including the Y chromosome gene RPS4Y1**

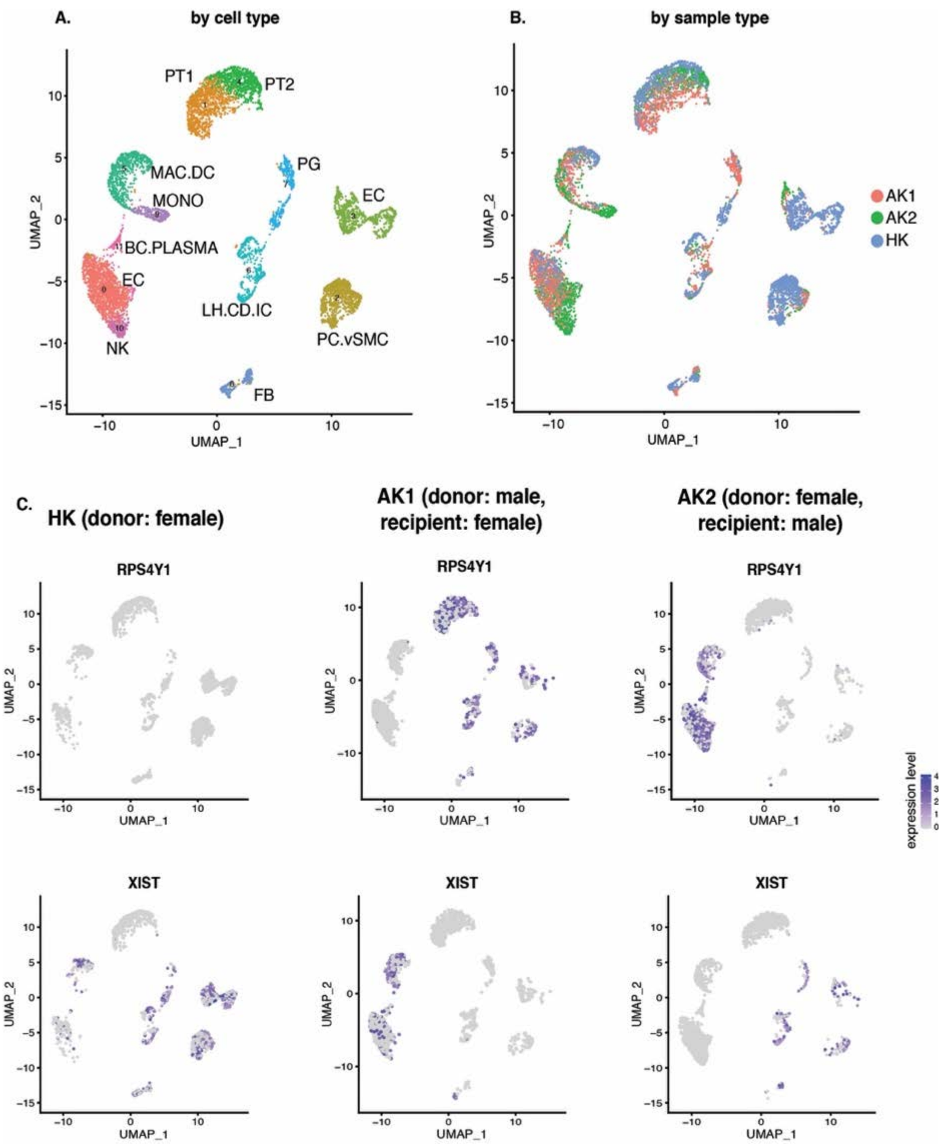


Figure depicts the UMAP-based visualization of 7217 individual cells obtained from the three kidney biopsy tissues shown in Figure 2 but including the Y-chromosome gene RPS4Y1. *EIF1AY, DDX3Y* (both shown in Figure 2) and *RPS4Y1* are Y chromosome-specific genes.

(A). Uniform manifold approximation and projection (UMAP)-based visualization by cell type.

(B). UMAP-based visualization by sample type.

(C). Feature plot showing expression of the Y-chromosome gene *RPS4Y1* (top panel) and X-chromosome gene *XIST* (bottom panel) in the three biopsies.
