## Supplementary material for "Detection of infiltrating fibroblasts by single-cell transcriptomics in human kidney allografts": S4 fig

### S4 Fig. Prediction scores for cell clusters when mapped to a kidney reference dataset

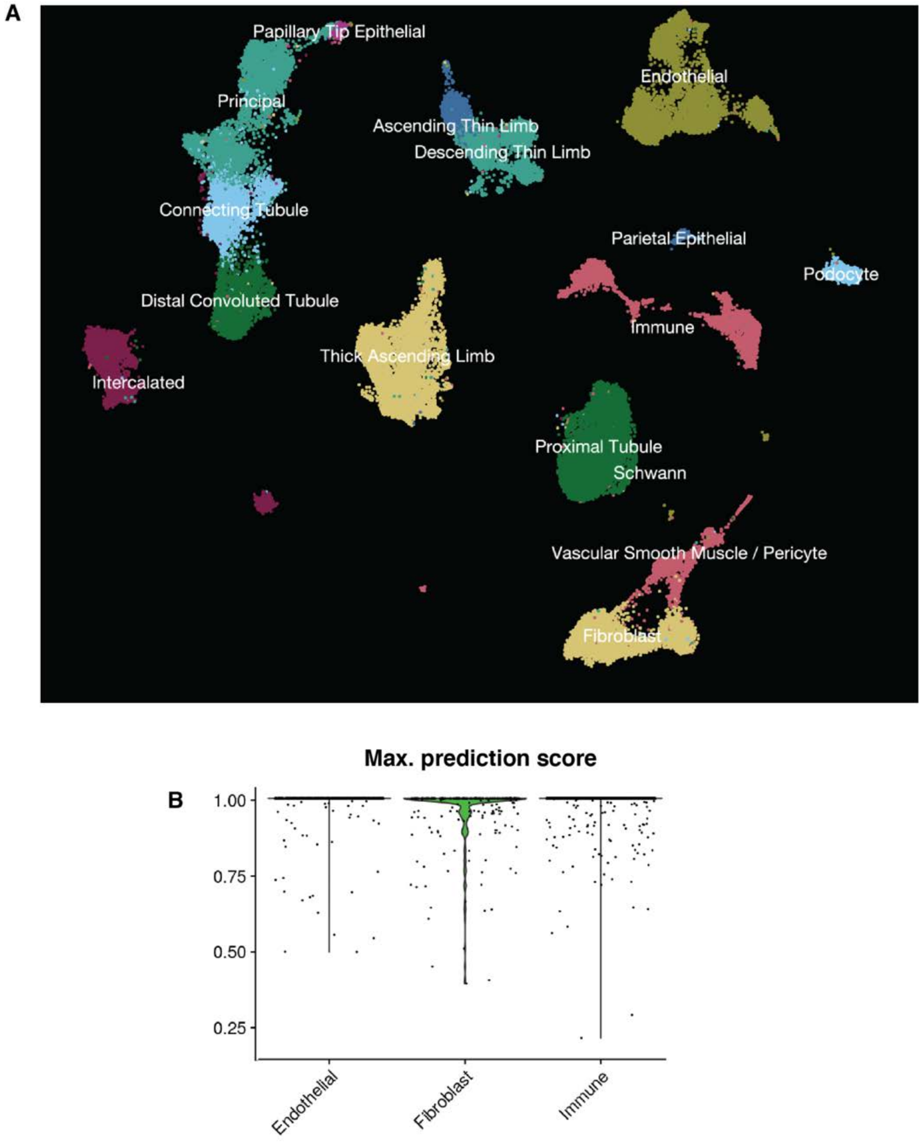


(A). Reference dataset consisting of 64,693 kidney cells generated in the Human Biomolecular Atlas Program (HuBMAP) and the Kidney Precision Medicine Project (KPMP) (25). The reference dataset represents 21 samples across 13 patient donors.

(B). We used ‘Azimuth’(23) to map our data to the annotated reference dataset. Most of the cells from our dataset mapped with high prediction score to the reference data; noticeably, as shown in the violin plot, fibroblasts mapped with very high prediction score, confirming the authenticity of our original annotation of fibroblasts.
