## Supplementary material for "Detection of infiltrating fibroblasts by single-cell transcriptomics in human kidney allografts": S5 fig

### S5 Fig. Agnostic mapping of cells from our study to the kidney reference dataset

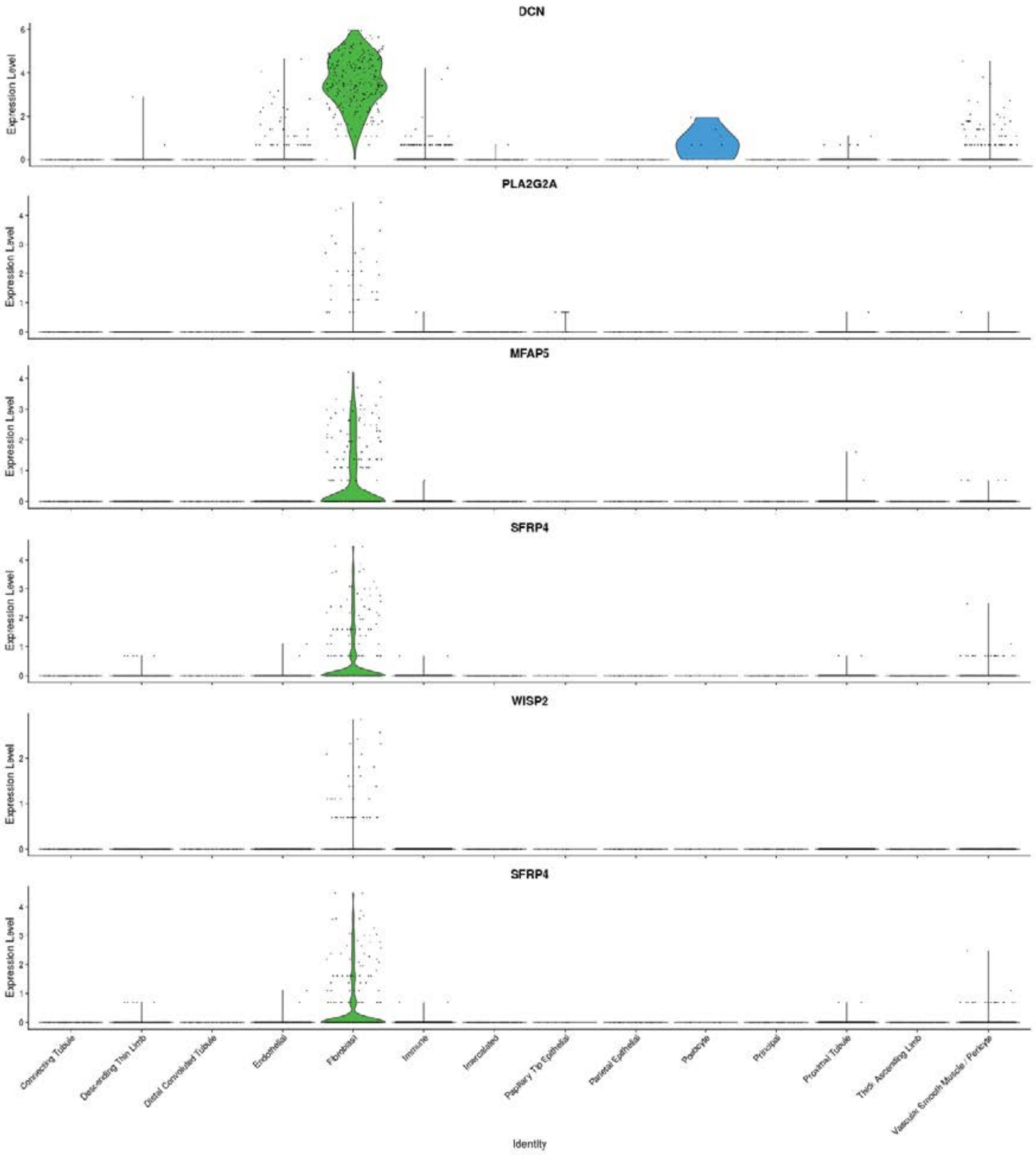


Single cell data from our study was agnostically mapped to a large-scale, publicly available, annotated, kidney reference dataset consisting of 64,693 kidney cells generated in the Human Biomolecular Atlas Program (HuBMAP) and the Kidney Precision Medicine Project (KPMP) (25). The reference dataset represents 21 samples across 13 patient donors. The violin plots show expression of genes, that corresponded to fibroblasts in our independent analysis, corresponds to fibroblast annotation in the reference dataset, thereby confirming accuracy of our annotation.
