## Supplementary material for "Detection of infiltrating fibroblasts by single-cell transcriptomics in human kidney allografts": S6 fig

### S6 Fig. Comparison of kidney fibroblasts with skin fibroblasts

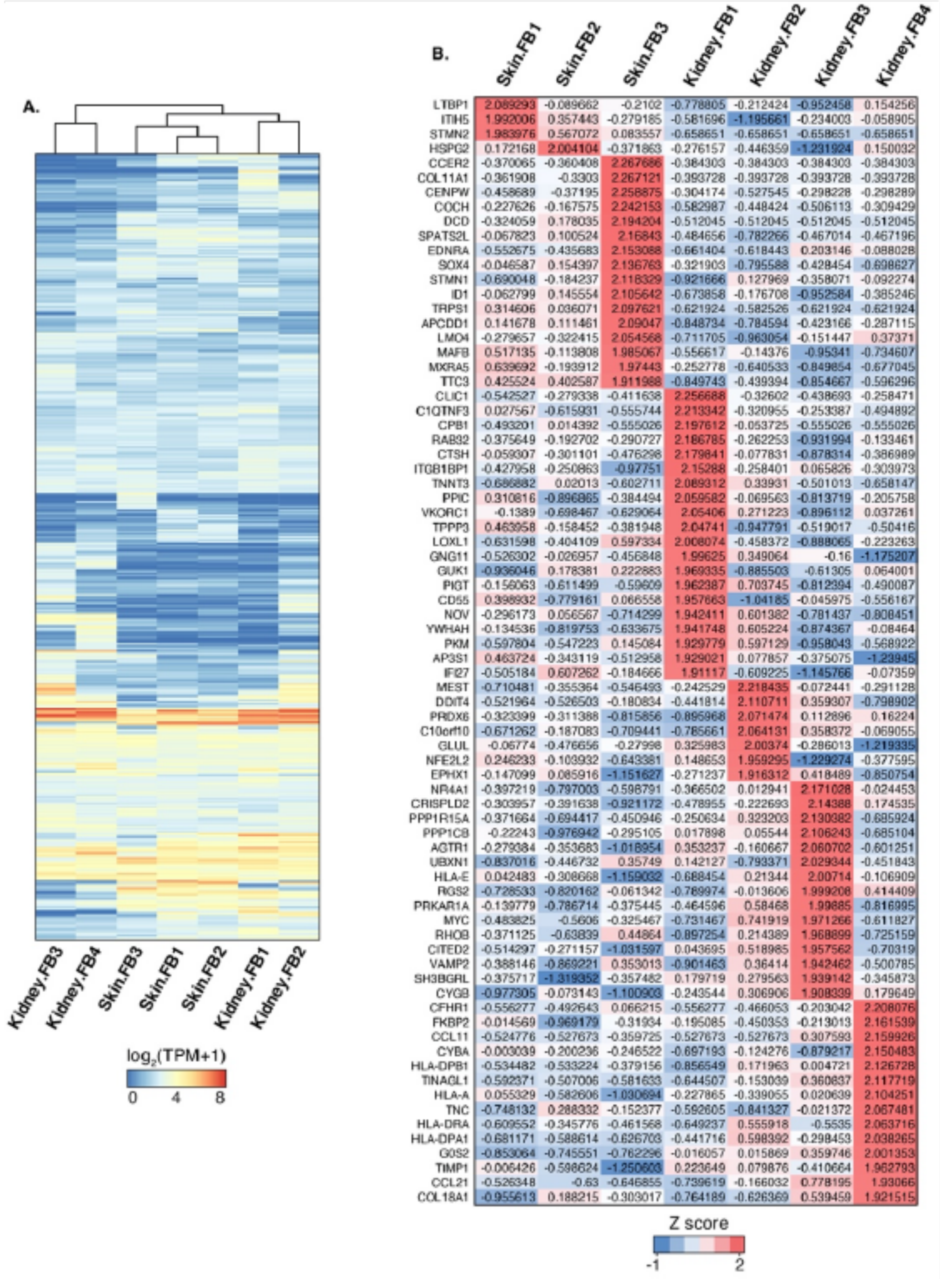


| | Skin.FB1 | Skin.FB2 | Skin.FB3 | Kidney.FB1 | Kidney.FB2 | Kidney.FB3 | Kidney.FB4 |
|---|---|---|---|---|---|---|---|
| LTBP1 | 2.089293 | -0.089662 | -0.2102 | -0.778805 | -0.212424 | -0.952458 | 0.154256 |
| ITIH5 | 1.992006 | 0.357443 | -0.279185 | -0.581696 | -1.195661 | -0.234003 | -0.058905 |
| STMN2 | 1.983976 | 0.567072 | 0.083557 | -0.658651 | -0.658651 | -0.658651 | -0.658651 |
| HSPG2 | 0.172168 | 2.004104 | -0.371863 | -0.276157 | -0.446359 | -1.231924 | 0.150032 |
| CCER2 | -0.370065 | -0.360408 | 2.267686 | -0.384303 | -0.384303 | -0.384303 | -0.384303 |
| COL11A1 | -0.361908 | -0.3303 | 2.267121 | -0.393728 | -0.393728 | -0.393728 | -0.393728 |
| CENPW | -0.458689 | -0.37195 | 2.258875 | -0.304174 | -0.527545 | -0.298228 | -0.298289 |
| COCH | -0.227626 | -0.167575 | 2.242153 | -0.582987 | -0.448424 | -0.506113 | -0.309429 |
| DCD | -0.324059 | 0.178035 | 2.194204 | -0.512045 | -0.512045 | -0.512045 | -0.512045 |
| SPATS2L | -0.067823 | 0.100524 | 2.16843 | -0.484656 | -0.782266 | -0.467014 | -0.467196 |
| EDNRA | -0.552675 | -0.435683 | 2.153088 | -0.661404 | -0.618443 | 0.203146 | -0.088028 |
| SOX4 | -0.046587 | 0.154397 | 2.136763 | -0.321903 | -0.795588 | -0.428454 | -0.698627 |
| STMN1 | -0.690048 | -0.184237 | 2.118329 | -0.921666 | 0.127969 | -0.358071 | -0.092274 |
| ID1 | -0.062799 | 0.145554 | 2.105642 | -0.673858 | -0.176708 | -0.952584 | -0.385246 |
| TRPS1 | 0.314606 | 0.036071 | 2.097621 | -0.621924 | -0.582526 | -0.621924 | -0.621924 |
| APCDD1 | 0.141678 | 0.111461 | 2.09047 | -0.848734 | -0.784594 | -0.423166 | -0.287115 |
| LMO4 | -0.279657 | -0.322415 | 2.054568 | -0.711705 | -0.963054 | -0.151447 | 0.37371 |
| MAFB | 0.517135 | -0.113808 | 1.985067 | -0.556617 | -0.14376 | -0.95341 | -0.734607 |
| MXRA5 | 0.639692 | -0.193912 | 1.97443 | -0.252778 | -0.640533 | -0.849854 | -0.677045 |
| TTC3 | 0.425524 | 0.402587 | 1.911988 | -0.849743 | -0.439394 | -0.854667 | -0.596296 |
| CLIC1 | -0.542527 | -0.279338 | -0.411638 | 2.256688 | -0.32602 | -0.438693 | -0.258471 |
| C1QTNF3 | 0.027567 | -0.615931 | -0.555744 | 2.213342 | -0.320955 | -0.253387 | -0.494892 |
| CPB1 | -0.493201 | 0.014392 | -0.555026 | 2.197612 | -0.053725 | -0.555026 | -0.555026 |
| RAB32 | -0.375649 | -0.192702 | -0.290727 | 2.186785 | -0.262253 | -0.931994 | -0.133461 |
| CTSH | -0.059307 | -0.301101 | -0.476298 | 2.179841 | -0.077831 | -0.878314 | -0.386989 |
| ITGB1BP1 | -0.427958 | -0.250863 | -0.97751 | 2.15288 | -0.258401 | 0.065826 | -0.303973 |
| TNNT3 | -0.686882 | 0.02013 | -0.602711 | 2.089312 | 0.33931 | -0.501013 | -0.658147 |
| PPIC | 0.310816 | -0.896865 | -0.384494 | 2.059582 | -0.069563 | -0.813719 | -0.205758 |
| VKORC1 | -0.1389 | -0.698467 | -0.629064 | 2.05406 | 0.271223 | -0.896112 | 0.037261 |
| TPPP3 | 0.463958 | -0.158452 | -0.381948 | 2.04741 | -0.947791 | -0.519017 | -0.50416 |
| LOXL1 | -0.631598 | -0.404109 | 0.597334 | 2.008074 | -0.458372 | -0.888065 | -0.223263 |
| GNG11 | -0.526302 | -0.026957 | -0.456848 | 1.99625 | 0.349064 | -0.16 | -1.175207 |
| GUK1 | -0.936046 | 0.178381 | 0.222883 | 1.969335 | -0.885503 | -0.61305 | 0.064001 |
| PIGT | -0.156063 | -0.611499 | -0.59609 | 1.962387 | 0.703745 | -0.812394 | -0.490087 |
| CD55 | 0.398932 | -0.779161 | 0.066558 | 1.957663 | -1.04185 | -0.045975 | -0.556167 |
| NOV | -0.296173 | 0.056567 | -0.714299 | 1.942411 | 0.601382 | -0.781437 | -0.808451 |
| YWHAH | -0.134536 | -0.819753 | -0.633675 | 1.941748 | 0.605224 | -0.874367 | -0.08464 |
| PKM | -0.597804 | -0.547223 | 0.145084 | 1.929779 | 0.597129 | -0.958043 | -0.568922 |
| AP3S1 | 0.463724 | -0.343119 | -0.512958 | 1.929021 | 0.077857 | -0.375075 | -1.23945 |
| IFI27 | -0.505184 | 0.607262 | -0.184666 | 1.91117 | -0.609225 | -1.145766 | -0.07359 |
| MEST | -0.710481 | -0.355364 | -0.546493 | -0.242529 | 2.218435 | -0.072441 | -0.291128 |
| DDIT4 | -0.521964 | -0.526503 | -0.180834 | -0.441814 | 2.110711 | 0.359307 | -0.798902 |
| PRDX6 | -0.323399 | -0.311388 | -0.815856 | -0.895968 | 2.071474 | 0.112896 | 0.16224 |
| C10orf10 | -0.671262 | -0.187083 | -0.709441 | -0.785661 | 2.064131 | 0.358372 | -0.069055 |
| GLUL | -0.06774 | -0.476656 | -0.27998 | 0.325983 | 2.00374 | -0.286013 | -1.219335 |
| NFE2L2 | 0.246233 | -0.103932 | -0.643381 | 0.148653 | 1.959295 | -1.229274 | -0.377595 |
| EPHX1 | -0.147099 | 0.085916 | -1.151627 | -0.271237 | 1.916312 | 0.418489 | -0.850754 |
| NR4A1 | -0.397219 | -0.797003 | -0.598791 | -0.366502 | 0.012941 | 2.171028 | -0.024453 |
| CRISPLD2 | -0.303957 | -0.391638 | -0.921172 | -0.478955 | -0.222693 | 2.14388 | 0.174535 |
| PPP1R15A | -0.371664 | -0.694417 | -0.450946 | -0.250634 | 0.323203 | 2.130382 | -0.685924 |
| PPP1CB | -0.22243 | -0.976942 | -0.295105 | 0.017898 | 0.05544 | 2.106243 | -0.685104 |
| AGTR1 | -0.279384 | -0.353683 | -1.018954 | 0.353237 | -0.160667 | 2.060702 | -0.601251 |
| UBXN1 | -0.837016 | -0.446732 | 0.35749 | 0.142127 | -0.793371 | 2.029344 | -0.451843 |
| HLA-E | 0.042483 | -0.308668 | -1.159032 | -0.688454 | 0.21344 | 2.00714 | -0.106909 |
| RGS2 | -0.728533 | -0.820162 | -0.061342 | -0.789974 | -0.013606 | 1.999208 | 0.414409 |
| PRKAR1A | -0.139779 | -0.786714 | -0.375445 | -0.464596 | 0.58468 | 1.99885 | -0.816995 |
| MYC | -0.483825 | -0.5606 | -0.325467 | -0.731467 | 0.741919 | 1.971266 | -0.611827 |
| RHOB | -0.371125 | -0.63839 | 0.44864 | -0.897254 | 0.214389 | 1.968899 | -0.725159 |
| CITED2 | -0.514297 | -0.271157 | -1.031597 | 0.043695 | 0.518985 | 1.957562 | -0.70319 |
| VAMP2 | -0.388146 | -0.869221 | 0.353013 | -0.901463 | 0.36414 | 1.942462 | -0.500785 |
| SH3BGRL | -0.375717 | -1.319352 | -0.357482 | 0.179719 | 0.279563 | 1.939142 | -0.345873 |
| CYGB | -0.977305 | -0.073143 | -1.100903 | -0.243544 | 0.306906 | 1.908339 | 0.179649 |
| CFHR1 | -0.556277 | -0.492643 | 0.066215 | -0.556277 | -0.466053 | -0.203042 | 2.208076 |
| FKBP2 | -0.014569 | -0.969179 | -0.31934 | -0.195085 | -0.450353 | -0.213013 | 2.161539 |
| CCL11 | -0.524776 | -0.527673 | -0.359725 | -0.527673 | -0.527673 | 0.307593 | 2.159926 |
| CYBA | -0.003039 | -0.200236 | -0.246522 | -0.697193 | -0.124276 | -0.879217 | 2.150483 |
| HLA-DPB1 | -0.534482 | -0.533224 | -0.379156 | -0.856549 | 0.171963 | 0.004721 | 2.126728 |
| TINAGL1 | -0.592371 | -0.507006 | -0.581633 | -0.644507 | -0.153039 | 0.360837 | 2.117719 |
| HLA-A | 0.055329 | -0.582606 | -1.030694 | -0.227865 | -0.339055 | 0.020639 | 2.104251 |
| TNC | -0.748132 | 0.288332 | -0.152377 | -0.592605 | -0.841327 | -0.021372 | 2.067481 |
| HLA-DRA | -0.609552 | -0.345776 | -0.461568 | -0.649237 | 0.555918 | -0.5535 | 2.063716 |
| HLA-DPA1 | -0.681171 | -0.588614 | -0.626703 | -0.441716 | 0.598392 | -0.298453 | 2.038265 |
| G0S2 | -0.853064 | -0.745551 | -0.762296 | -0.016057 | 0.015869 | 0.359746 | 2.001353 |
| TIMP1 | -0.006426 | -0.598624 | -1.250603 | 0.223649 | 0.079876 | -0.410664 | 1.962793 |
| CCL21 | -0.526348 | -0.63 | -0.646855 | -0.739619 | -0.166032 | 0.778195 | 1.93066 |
| COL18A1 | -0.955613 | 0.188215 | -0.303017 | -0.764189 | -0.626369 | 0.539459 | 1.921515 |

We compared the gene expression profiles of kidney fibroblasts with that of skin fibroblasts. We have reported 3 fibroblast subtypes by scRNA-seq of human skin from 5 patients with atopic dermatitis and 7 healthy control subjects *(He et al. J Allergy Clin Immunol 2020 PMID:* ***32035984).*** In order to compare the expression profile of FBs in the allograft kidney with the FBs in skin, we selected union of top 100 genes expressed in each FB population and performed unsupervised clustering as seen in the heatmap (Panel A). Next, we analyzed the most abundant and differentially expressed genes in all the FBs using the 'Z score' (Panel B).
